# Stars2Cells: Astrometric Tracking of Neurons Across Imaging Sessions

**DOI:** 10.64898/2026.07.03.736144

**Authors:** Ari Peden-Asarch, Lauren Honan, Jacqueline Bai, Eli Asarch, Josiah Quinn, Kevin R. Coffey, John F. Neumaier

## Abstract

Chronic calcium imaging offers a window into how single neurons and ensemble activity change across days where identifying the same neurons from one session to the next is the prerequisite for answering questions regarding learning, drift, and plasticity over time. Yet only ∼2–3% of imaging laboratories publish longitudinal cross-session work, because existing registration tools depend on spatial-footprint or temporal correlations that degrade under repeated recording sessions. Here, we introduce **Stars2Cells (S2C)**, a tracking pipeline inspired by astrometric plate-solving that represents each neuron’s local geometry as a four-dimensional quad descriptor invariant to rotation, translation, and uniform scaling. S2C operates purely on centroid coordinates and combines descriptor-space matching, Random Sample Consensus (RANSAC) verification, and Hungarian assignment. Across a synthetic benchmark of 1,262 paired runs spanning 100–1,000 neurons and 8 perturbation conditions plus 1 identity sanity-floor, S2C reached pooled F1 = 98.4% compared to the standard ROI-based matching of 36.0%. To show what this enables, we applied the pipeline to dorsomedial striatum (DMS) imaging during oral fentanyl behavioral-economics self-administration. Here, we show that a conserved population-rewarded lever press response in DMS masks near-complete single-neuron turnover. This representational-drift signature we demonstrated is invisible to the bulk photometry, and resolving it requires the same-cell tracking S2C provides. S2C is distributed as a GUI-driven standalone application for both macOS and Windows, requiring no Python, command line, or virtual environment setup.

## 1 Introduction

Chronic two-photon microscopy through cranial windows first established that the same cortical neurons could be revisited over days and weeks, enabling imaging-based single-cell investigations of learning, plasticity, and representational drift *in vivo* [1–3]. Miniaturized one-photon fluorescence microscopes (“miniscopes”) extended this access to freely behaving animals and to deep-brain structures inaccessible through head-fixed objectives [4–8]. Regardless, re-identifying a given neuron from one session to the next is the prerequisite for any longitudinal investigation, and the difficulty in reliably identifying the same neurons from session to session has prevented researchers from taking full advantage of recurrent imaging.

To quantify this disparity, we surveyed PubMed, where two-photon and miniscope publications have grown steeply over the past decade (Fig. 1a). Combined, *∼*8,484 unique last authors have published two-photon calcium-imaging work (with a further *∼*500 UCLA Miniscope and *∼*600 Inscopix laboratories estimated from manufacturer/community figures; Fig. 1b), yet only *∼*220 (*∼*2.3%) unique last authors have published longitudinal cross-session imaging since 2021. Tool-level adoption is still sparser with citation counts of each tool’s reference publication (OpenAlex, accessed 06/2026) as follows: CellReg [9] leads with 435 citations, reflecting both an eight-year head start and the footprint-correlation family’s incumbency in 1P miniscope data. SCOUT [10] follows with 12 citations and “Track2p” [11] with 1 citation, spanning longitudinal-pipeline and developmental-2P use, respectively; CaliAli [12] has 4 citations; both were published within the past year, and thus still below the citation horizon. Among cross-session image-registration tools, PatchWarp [13] has 29 citations and the fully affinity-invariant FOV-alignment method FAIM [14] has 3 citations; and EMC^2^ [15], for continuous tracking within the session, has 33 citations.

**Figure 1:**
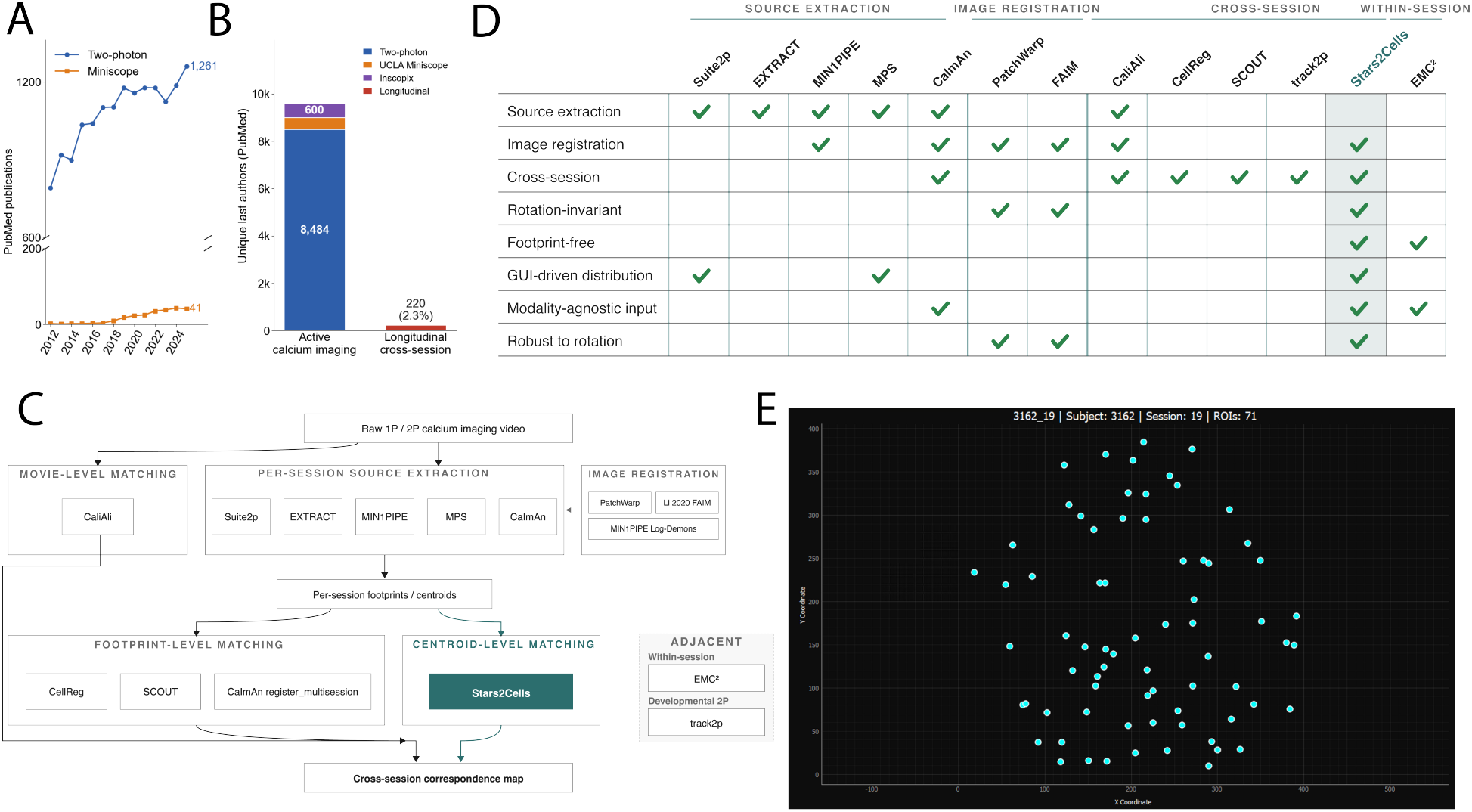
The longitudinal tracking gap, existing tool landscape, and the GUI-driven S2C application. **a**, PubMed publications per year for “two-photon” (blue) and “miniscope” (orange) (2012–2025), showing rapid growth in active calcium imaging. **b**, Active imaging labs versus labs publishing longitudinal cross-session tracking. **c**, Feature comparison of the calcium-imaging analysis tool landscape. **d**, Layer diagram of the cross-session analysis pipeline. **e**, Stars2Cells GUI-driven standalone application.

These tools span three layers of the pipeline, distinguished by the input each consumes (Fig. 1c): *movie-level* methods (CaliAli [12]) register raw video on landmarks such as vasculature before extraction; *footprint-level* methods (CellReg [9], CaImAn’s register_multisession [16], SCOUT [10]) match per-session ROI masks by spatial correlation; and *centroid-level* matching, where S2C operates, recovers correspondence directly from the 2D point geometry any extractor produces. Each layer trades one cue for another (Fig. 1d): movie-level needs visible vasculature, footprint-level needs ROI shape stability across days, and centroid-level methods discard both but runs on any extractor’s output across any modality (1P, 2P, mesoscope, hand-drawn), handles rotation by construction, and applies to archived data. S2C delivers this as a GUI-driven standalone application for macOS and Windows (Fig. 1e) that carries raw recordings through to multi-session neuron tracks end-to-end.

Two converging bottlenecks have kept this longitudinal imaging out of the scientific zeitgeist. First, raw single-photon miniscope video processing historically required deep computational expertise. Pipelines such as CaImAn and Minian asked end users to assemble Python or MATLAB environments, configure GPU/parallel backends, manually tune CNMF/CNMF-E parameters, and inspect results from the command line, which alone can take new groups weeks to months to operationalize [16, 17]. MPS [18] addresses this upstream side by consolidating preprocessing, motion correction, source extraction, deconvolution, and quality control behind a single graphical interface, removing the engineering overhead that historically gated entry for non-engineering groups. Second, reliably matching the same cells across recording days has remained unsolved at the level of inter-session FOV variability that single-photon recordings regularly produce. The incumbent approach matches cells by correlating their per-session ROI footprints (hereafter *ROI matching*), the strategy behind CellReg [9] and the footprint-correlation methods that followed it. It assumes tightly co-registered fields of view and stable footprint shape across days, both of which break under the baseplate-reattachment rotations, translational drift, and detection dropout that chronic single-photon recordings routinely produce. However, this disparity is not driven by lack of interest in longitudinal questions, but reflects the practical difficulty of reliably matching neurons across sessions with existing tools, layered on top of the overhead of installing, configuring, and operating Python-based pipelines.

The connection to astronomy is the idea that first prompted this project: how does a telescope point to an unfamiliar patch of sky and know where in space it is? The answer is astrometric plate-solving, which is the routine, well-validated procedure by which astrometric pipelines such as Astropy and Astrometry.net match an unannotated star field to a catalog despite unknown rigid transformations [19, 20]. The solution emerges from local geometry where each star is described by the geometry of its four-star neighborhood (a quad), and the resulting normalized descriptor is invariant to rotation, translation, and uniform scaling [19, 20]. Matching is performed in descriptor space, followed by geometric verification via RANSAC, yielding robust correspondences even when large fractions of the field are obscured. However, this machinery only solves *where* a telescope is pointing, and thus, does not assign astronomical identities to individual stars. Therefore, S2C is the natural followup intended for the neuron-tracking problem across sessions and/or days.

The centroid coordinates produced by MPS [18], Suite2p [21], CaImAn [16], Minian [17], MIN1PIPE [22], EXTRACT [23], or hand-drawn ROIs all naturally feed into S2C. Given our lab produced MPS [18], we designed MPS and S2C to form a complete GUI-driven workflow from raw miniscope video to longitudinal neuron tracking, distributed as GUI-driven standalone applications for both macOS and Windows (Fig. 1e). The pipeline proceeds in five automated steps: (1) quad descriptor generation from centroid constellations using a diagonal-first algorithm with KNN local and random long-range sampling; (2) descriptor-space matching with an automatically calibrated similarity threshold that scales as 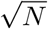, where *N* is the number of neurons; (3) RANSAC geometric verification to reject false-positive descriptor matches; (4) Hungarian assignment to produce globally optimal one-to-one neuron correspondences; and (5) consolidation of pairwise matches into multi-session neuron tracks. Each step is visualized and inspectable through an integrated GUI; each build bundles a self-contained Python runtime and all dependencies (a signed and notarized DMG for macOS, a portable executable for Windows), so no manual dependency configuration is required.

The dorsomedial striatum (DMS) is the canonical associative striatal hub for action-outcome learning, value computation, and goal-directed control [24, 25], and dorsal/dorsomedial striatum is causally implicated in relapse to opioid seeking after voluntary abstinence [26], with dorsal-striatal connectivity tracking the incubation of opioid craving [27]. DMS dopamine signaling longitudinally tracks the emergence of compulsive reward seeking across self-administration [28]. Yet despite this causal centrality, to our knowledge every published *in vivo* recording in dorsomedial striatum *cell bodies* during *opioid* self-administration to date has been bulk neurochemical (fiber photometry) via dopamine, glutamate, or aggregated Ca^2+^ activity [28–31] leaving the cellular composition and longitudinal evolution of DMS dynamics during opioid taking uncharacterized at single-cell resolution. How DMS output neurons reorganize across chronic opioid exposure has been characterized almost exclusively *ex vivo*, and *in vivo* only in nucleus accumbens via miniscopes [32–35]. In parallel, behavioral-economics demand procedures [36, 37] yield scale-invariant, orthogonal measures of intake (*Q*_0_) and motivation (*α*) that uniquely predict compulsivity, reinstatement, and pharmacotherapy efficacy, including for fentanyl [30, 38, 39]. Yet these measures have been combined only with bulk neurochemical signals and not with single-cell activity. Thus, since it has been established across hippocampus, cortex, and striatum population-level signals can mask near-complete turnover of the underlying ensemble, then asking whether the same DMS neurons carry the press response from session to session across an escalating fentanyl demand curve is impossible without same-cell tracking. The MPS→S2C stack applied here provides the first cell-resolved longitudinal recording of individually tracked dorsomedial striatum neurons across an escalating opioid demand curve, to our knowledge.

The website for download is available at https://ariasarch.github.io/Stars2Cells/, a sample walkthrough at https://github.com/ariasarch/Stars2Cells_Sample_Code, and the source code at https://github.com/ariasarch/Stars2Cells_GUI.

## 2 Results

### 2.1 Quad descriptor pairing from neuron centroid constellations

S2C represents each neuron’s spatial context by enumerating four-neuron patterns (quads) from the centroid field and computing a normalized geometric descriptor for each (Fig. 2a). Given *N* centroid positions in a session, the pipeline constructs candidate diagonals by combining *k*-nearest-neighbor local edges with random long-range edges, ensuring descriptors capture both fine local structure and FOV-spanning geometry (Fig. 2b). For each diagonal, perpendicular heights are computed for all other neurons, and the top-*K* third points by height are retained (default *K* = 25 –tunable). Third points are then paired to form quads, and a four-dimensional descriptor (*x*_*C*_, *y*_*C*_, *x*_*D*_, *y*_*D*_) is computed by projecting the two non-diagonal vertices onto the coordinate frame defined by the quad’s longest pairwise edge, normalizing by the diagonal length.

**Figure 2:**
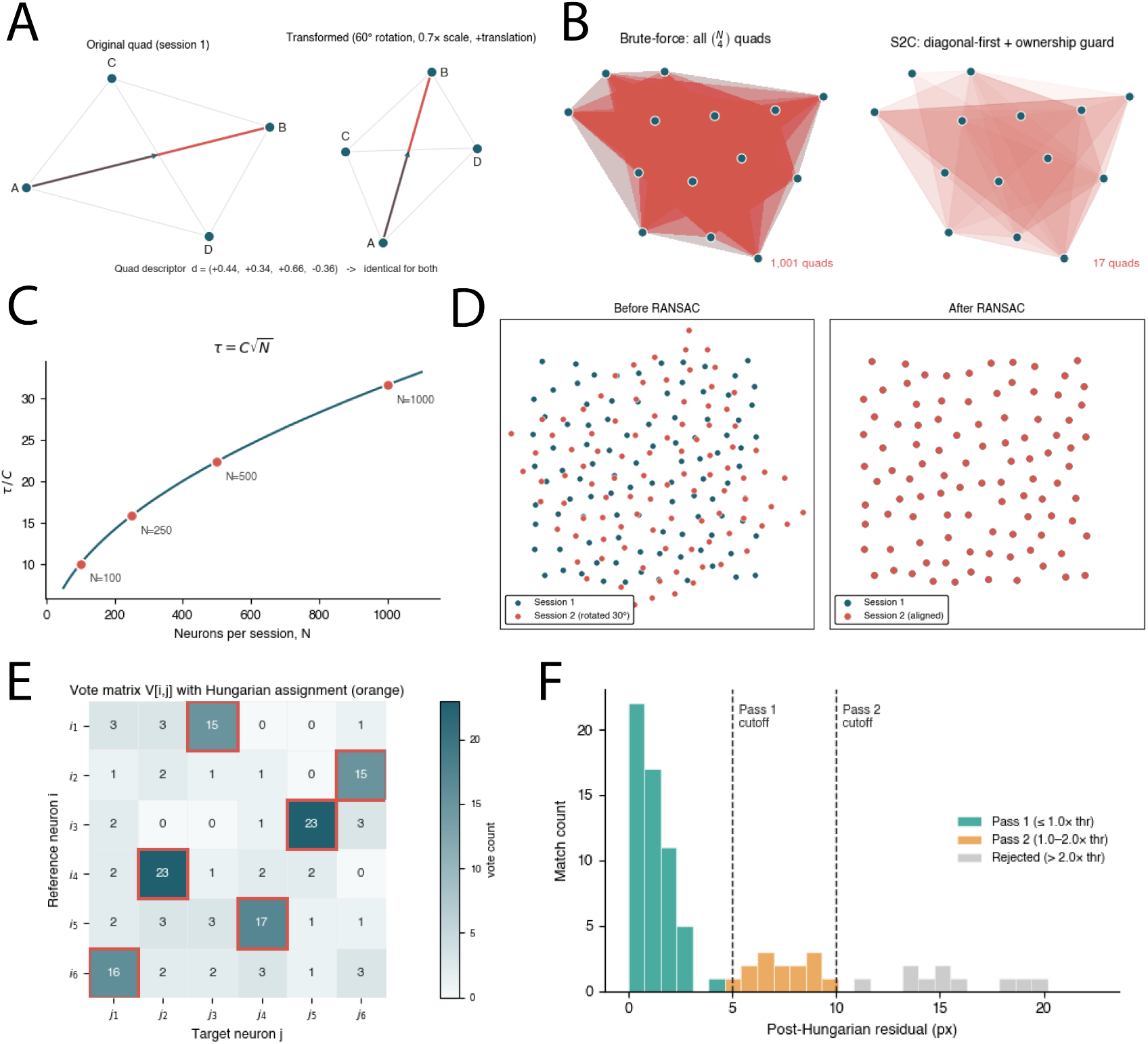
The Stars2Cells pipeline. **a**, Astrometric metaphor: a four-star quad’s relative geometry, normalized on its longest edge. **b**, Step 1, diagonal-first quad enumeration with ownership guard. Each translucent polygon spans the four neurons of one enumerated quad; cumulative compositing renders quad density as color saturation. *Left:* brute-force enumeration of all 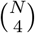 quads on an example field (*N* = 14 neurons; 1,001 quads) saturates into an unreadable red blob where no single quad is resolvable. *Right:* S2C’s diagonal-first construction (*k*_local_ = 15 KNN local diagonals plus *k*_random_ long-range partners per neuron at production scale; scaled down to *k*_local_ = 4, *k*_random_ = 2 here so the *N* = 14 stays readable) emits each unique quad exactly once from its longest pairwise edge, yielding a sparse selective set in which individual quads remain visible. Complexity: *O*(*NkK*^2^) vs. *O*(*N* ^4^) for brute force. **c**, Step 1.5, empirical similarity-threshold scaling: the optimal threshold *τ* scales as 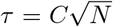 across the four benchmark neuron counts (*N* = 100, 250, 500, 1,000). **d**, Step 2.5, vectorized RANSAC: 100 simulated neurons (Poisson-disk-sampled in a 600*×* 600 FOV) with session 2 rotated 30^*•*^ about the FOV centre plus a small translation (left); after RANSAC fits the rigid transform from *K* = 1,000 batched-SVD candidate transforms and the inverse is applied, session 2 (orange) registers exactly to session 1 (teal) (right). **e**, Step 3, vote matrix and Hungarian assignment: each verified quad votes for 4 *×*4 = 16 neuron pairings; degree-normalized votes plus a spatial-residual term form the cost matrix, which is padded with rectangular dummy slots so unmatchable neurons can stay unmatched. **f**, Pass 1 versus Pass 2 recovery (residual distribution by pass): Pass 1 enforces a tight 1.0*×* RANSAC residual cutoff (green); Pass 2 re-solves a smaller Hungarian over the unmatched residual under a relaxed 2.0*×* cutoff (orange), recovering correspondences that Pass 1 conservatively rejected; matches with residual *>* 2.0*×* the RANSAC threshold are rejected outright (grey).

A key algorithmic innovation is the ownership guard: each quad is emitted exactly once, from the single diagonal equal to its longest pairwise edge (Methods; Fig. 2b). This avoids both the intractable 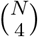 scaling of brute-force enumeration and the 6*×* duplicate emission of a guard-free construction.

Quad saturation analysis confirmed that the median nearest-neighbor distance in descriptor space (NN_*d*_ = 0.0101) exceeded the median descriptor blur (*σ*_*d*_ = 0.0041) for all sessions, with a saturation ratio of 2.46, indicating that the descriptor field was not saturated and individual neurons remained distinguishable.

Accurate neuron matching requires a larger similarity metric when neuron count grows. Thus, this is built into the next step of S2C via descriptor matching that requires a similarity threshold *τ* below which two quad descriptors are considered a match. As a field densifies, local-diagonal lengths shrink and the per-coordinate blur of a matched pair rises. Thus, since same-cell descriptor blur grows with neuron count, a fixed threshold rejects true matches at large *N* and admits false ones at small *N*, so *τ* must grow with field size. The empirical 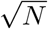 scaling is illustrated at the four benchmark neuron counts in Fig. 2c. S2C automates the per-animal calibration of *C* in Step 1.5 by sweeping thresholds on sampled quad subsets for each animal and fitting via least-squares regression through the origin. The resulting 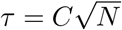 is applied per animal as the Step 2 descriptor threshold; it is a geometry-motivated estimate rather than a precisely calibrated value (Supplementary Note 2), with any residual miscalibration absorbed downstream by the consistency filter and RANSAC, so that registration accuracy is robust to its exact value (98.4% mean F1; Fig. 4).

Nonetheless, raw descriptor matches still contain both true correspondences and false positives from coincidental descriptor similarity. S2C applies a two-stage filtering process. First, a consistency filter removes matches whose implied spatial transformation is inconsistent with the consensus. Second, vectorized RANSAC [40] estimates the dominant rigid transformation between session pairs from matched quad centroid positions, retaining only geometrically consistent inlier matches (Fig. 2d).

The surviving quad-level matches are aggregated into a neuron-level vote matrix *V*, where *V* [*i, j*] counts how many verified quad matches jointly contain reference neuron *i* and target neuron *j*. The Hungarian algorithm [41] is applied to the cost matrix (max(*V* ) *− V* ) to produce globally optimal one-to-one neuron assignments. A cost-threshold sweep identifies the assignment threshold that maximizes matched neurons while preserving geometric coherence (Fig. 2e). A two-pass solve recovers correspondences that a single conservative cutoff would discard: Pass 1 enforces a tight 1.0*×* RANSAC-residual cutoff (tunable), Pass re-solves a smaller Hungarian over the unmatched residual under a relaxed 2.0*×* cutoff, and matches with residual exceeding 2.0*×* the RANSAC threshold (tunable) are rejected outright (Fig. 2f).

### 2.2 Activity correlation alone cannot identify neurons across sessions

A natural question is whether activity-based features such as cross-session calcium-trace correlation could substitute for or complement geometric matching. We bound its contribution from above with a deliberately aggressive idealized scenario that hands the temporal-correlation matcher every advantage geometry could provide: 1,000 simulated neurons placed at *identical* spatial positions across five sessions. Because positions are identical, any geometric or footprint-based matcher scores perfectly, so the test isolates the contribution of activity alone.

For each pair of sessions, we computed the full 1,000 *×* 1,000 Pearson correlation matrix between *z*-scored traces (in float32 row-chunks of 100 to control memory) and assigned each neuron in session *s*_1_ to the maximally correlated neuron in session *s*_2_. Performance was evaluated against ground-truth identity across recording durations of 10, 20, 60, and 180 min (6,000 to 108,000 frames per session). At every tested duration, mean matching accuracy across the 10 session pairs fell within the chance band: 0.12% [0.06, 0.19] at 10 min (*p* = 0.53 vs. chance), 0.14% [0.09, 0.20] at 20 min (*p* = 0.38), 0.07% [0.01, 0.14] at 60 min (*p* = 0.56), and 0.15% [0.08, 0.24] at 180 min (*p* = 0.53); chance is 1*/N* = 0.10% (Fig. 3). Thus, the activity-only matcher recovered no meaningful identity signal as each fell within the chance band at every duration.

**Figure 3:**
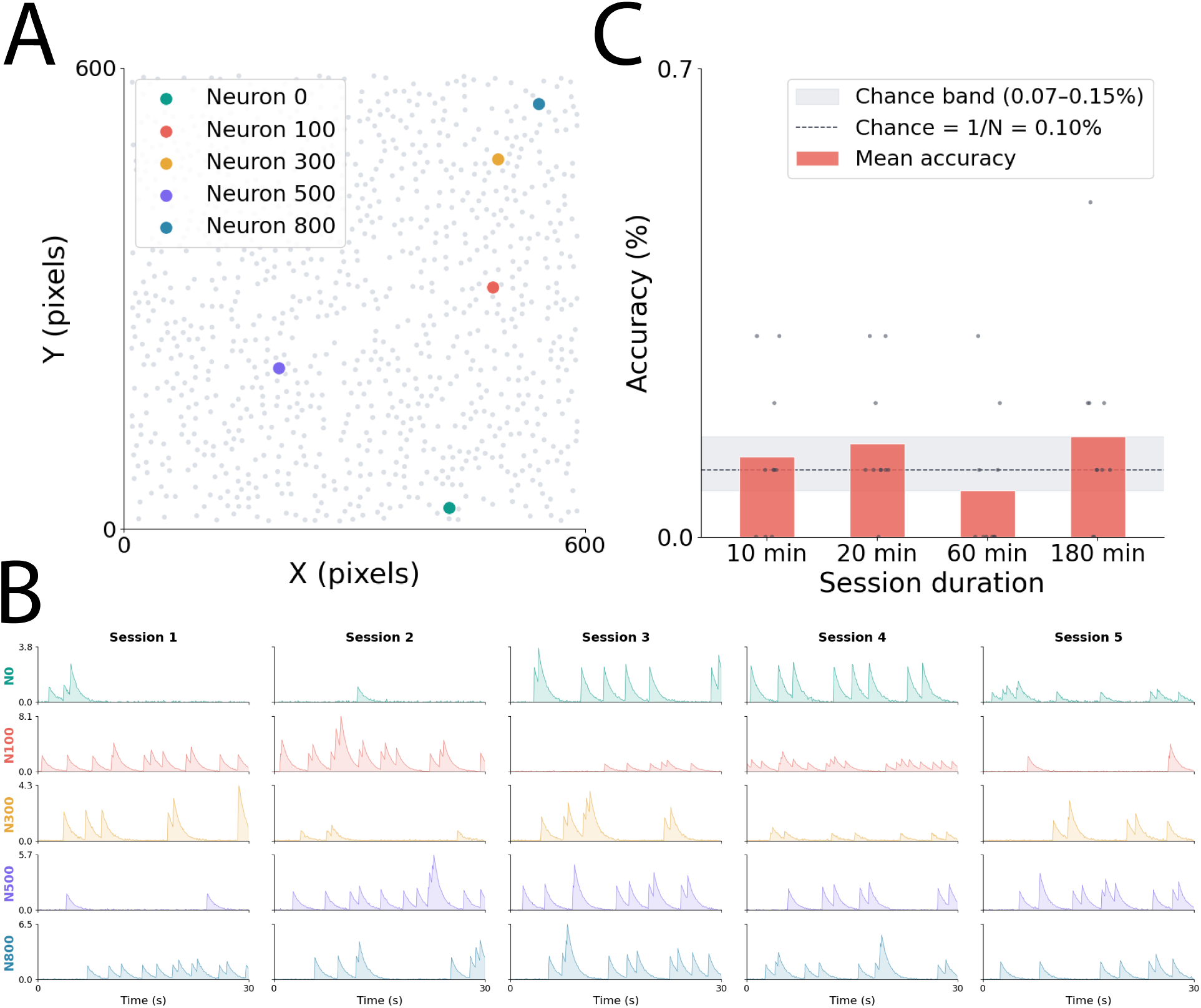
Temporal correlation alone is chance-level for cross-session matching. **a**, Spatial map: 1,000 simulated neurons placed by Poisson-disk sampling in a 600 *×* 600 px FOV; the same layout is reused across all 5 sessions, so a geometric matcher always would score 100%. Five example neurons are highlighted (colored) for panel **b. b**, Example AR(1) calcium traces (first 30 s shown) for the five highlighted neurons across sessions 1–5. Despite identical spatial identity, dynamics are statistically independent across sessions – each session reflects a fresh draw from that neuron’s generative distribution. **c**, Mean cross-session matching accuracy (mean *±*scatter across 10 session pairs) from a max-correlation matcher remains within the chance band (0.07–0.15%; chance 1*/N* = 0.10%, dashed line) at all four tested durations (10, 20, 60, 180 min); *p >* 0.3 vs. chance at every duration.

### 2.3 Synthetic validation across recording modalities

We validated S2C on a synthetic benchmark of 9 inter-session degradation conditions evaluated at four neuron-count tiers (100, 250, 500, 1,000), chosen to span the densities seen across calcium-imaging preparations: *∼*50–200 neurons per FOV in deep-brain miniscope recordings of DMS, habenula, dorsal raphe, and BNST [8]; *∼*200–500 in cortical and hippocampal miniscope recordings [5, 42]; and *∼*500–1,000+ in two-photon cortical imaging [21]. Degradation parameters were selected to bracket the inter-session perturbations we and others have observed in chronic miniscope recordings, while also extending into scenarios that the field’s hardware and surgical practice could plausibly produce: baseplate-reattachment rotation and animal head motion against the implant (uncorrected by MPS’s translational-only motion correction), residual translational drift downstream of MPS, ROI dropout from CNMF detection variability, and centroid localization noise. The ranges therefore combine direct empirical observation with a forward-looking margin for the more challenging operating conditions chronic experiments may encounter.

#### S2C is accurate and robust across the full benchmark parameter space

S2C achieved a mean F1 score of 98.4% across 1,262 paired runs spanning 8 conditions and 32 productive (condition *×* neuron-count) cells, with precision 99.9% and recall 97.2% (Fig. 4c). Performance was robust across neuron counts: F1 = 99.7% at *N* = 100, 99.9% at *N* = 250, 97.0% at *N* = 500, and 97.3% at *N* = 1,000 (Fig. 4d). Across tiers (illustrated at *N* = 250), S2C reached 99.9% F1 on Tier A (2P-like), 99.9% on Tier B (Miniscope-Like), and 99.9% on Tier C (Extreme) (Fig. 4e). Per-tier dips at higher neuron counts were driven by recall (precision remained near-saturated, *≥* 99.4% across all neuron counts), reflecting genuinely missed correspondences in the densest, most-perturbed conditions rather than spurious matches. Transitivity consistency across five chained sessions was high: pairwise F1 across the four consecutive transitions remained essentially flat at 98.3%, 98.4%, 98.4%, and 98.4% for A→B, B→C, C→D, and D→E respectively, indicating that registration accuracy is independent of position within the chain and does not degrade as sessions accumulate (Fig. 4f).

**Figure 4:**
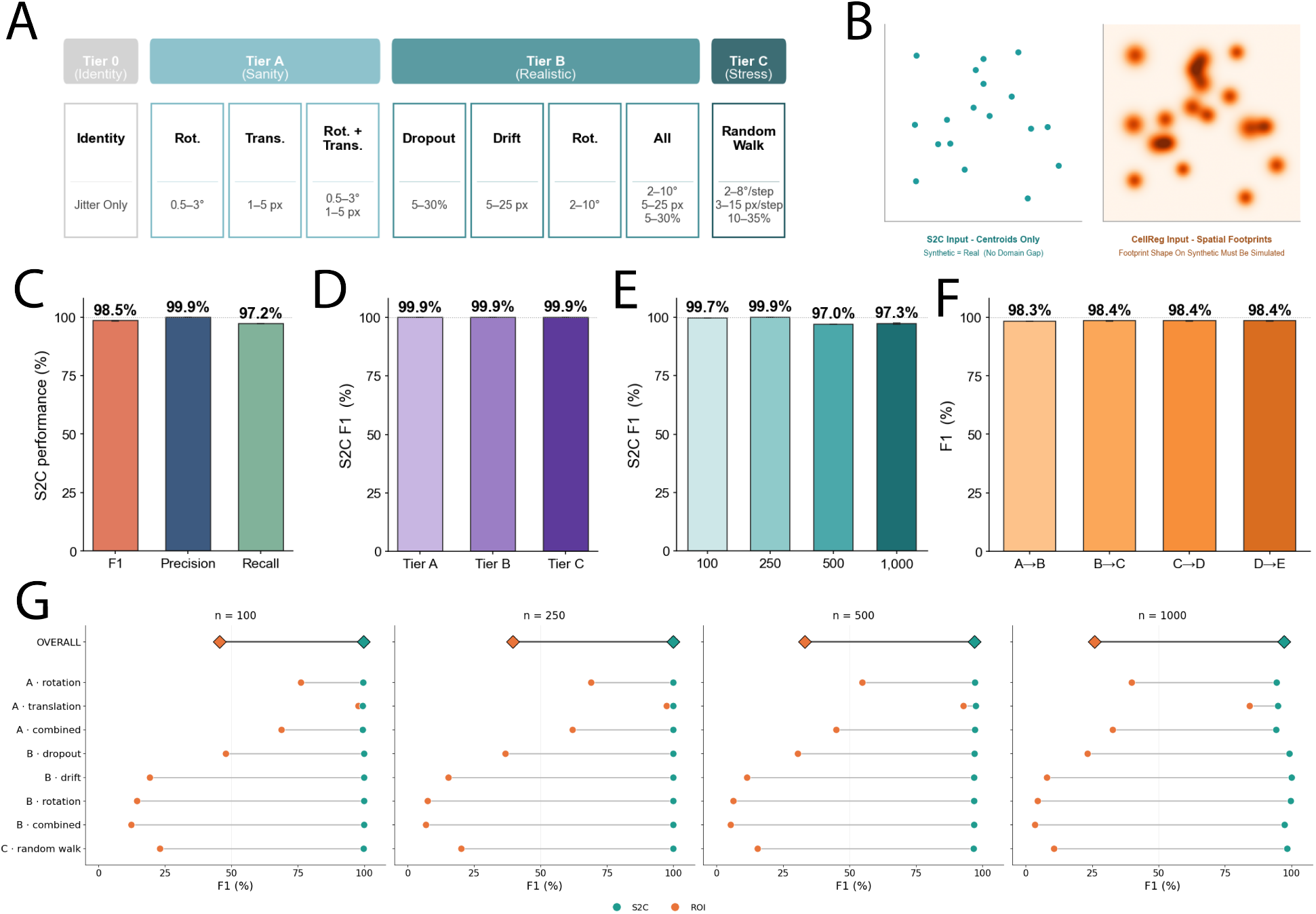
S2C is robust across the full benchmark parameter space and outperforms ROI-based matching at every condition. **a**, Benchmark structure: 8 productive conditions plus 1 identity sanity-floor (Tier 0 identity / Tier A (2P-Like) 3 / Tier B (Miniscope-Like) 4 / Tier C (Extreme) 1) *×* 4 neuron counts (100, 250, 500, 1,000) *×* 8 animals *×* 5 seeds *×* 5 consecutive sessions. **b**, Domain-gap-free validation: because S2C operates exclusively on 2D centroid coordinates, the input distribution is identical for synthetic and real recordings. ROI-based matching, in contrast, requires footprint shape that must be simulated approximately. **c**, Overall F1, precision, and recall pooled across all conditions and neuron counts. **d**, F1 stratified by neuron count. **e**, F1 stratified by tier at fixed *N* = 250. **f**, Per-transition F1 across five-session chains: pairwise F1 for the four consecutive transitions (A →B, B →C, C→ D, D →E). Performance is stable across chain position, with no per-step degradation. **g**, S2C versus ROI-based matching: dumbbell plot of paired F1 (S2C, teal •; ROI-based matching, orange• ) under the worst-case (perturbed) footprint scenario, one column per neuron count (100, 250, 500, 1,000). Each row is one of 8 degradation conditions (Tier-A rotation/translation/combined; Tier-B dropout/drift/rotation/combined; Tier-C random walk); diamond markers (**OVERALL**) pool all 8 conditions per neuron count (up to *n* = 320 paired runs each). S2C outperforms at every cell, with a pooled S2C *−≤*ROI-based matching gap of +54.1, +60.3, +63.9, and +71.4 pp at *N* = 100, 250, 500, 1,000. The ROI-based matching deficit is geometric (rotation/translation/dropout breaking the spatial-correlation signal upstream) rather than footprint-shape-driven: per-neuron-count ΔF1 is essentially identical between best-case and worst-case conditions (matched-condition surface, 100–500 neurons, all 8 conditions; |Δ_best-case_ *−*Δ_worst-case_| ≤≤0.3 pp). Grand-pooled paired mean ΔF1 = +62.4 pp [+60.5, +64.4], *d*_*z*_ = +1.81, *p <* 0.001 across 1,262 paired runs (per-condition: paired Wilcoxon signed-rank, Holm-corrected; effect sizes paired Cohen’s *d*_*z*_; 95% CIs bias-corrected bootstrap).

#### S2C achieved higher F1 than ROI-based matching at every neuron count and on every condition tested

Pooling within each neuron-count tier under the worst-case footprint scenario (complete shape perturbation; Fig. 4g): at *N* = 100, S2C reached F1 = 99.7% vs. ROI-based matching’s 45.6% (ΔF1 = +54.1 pp, 95% CI [+50.0, +58.1], paired Cohen’s *d*_*z*_ = +1.49, *p* = 1.1 *×* 10^*−*51^ Holm-corrected); at *N* = 250, F1 = 99.9% vs. 39.6% (Δ = +60.3 pp [+56.2, +64.3], *d*_*z*_ = +1.67, *p* = 1.8 *×* 10^*−*53^); at *N* = 500, F1 = 97.0% vs. 33.1% (Δ = +63.9 pp [+60.3, +67.5], *d*_*z*_ = +1.98, *p* = 3.3 *×* 10^*−*52^); at *N* = 1,000, F1 = 97.3% vs. 25.9% (Δ = +71.4 pp [+67.9, +74.6], *d*_*z*_ = +2.34, *p* = 3.5 *×* 10^*−*53^). Pooled across all 1,262 paired runs, S2C reached F1 = 98.4% vs. ROI-based matching’s 36.0% (paired mean ΔF1 = +62.4 pp [+60.5, +64.4], *d*_*z*_ = +1.81, *p <* 0.001; Fig. 4g, dumbbell panels per neuron count). Restricting the pool to the empirically-grounded conditions (Tiers A and B) demonstrates that the pooled gap is not an artifact of the most extreme conditions: S2C F1 = 98.4% vs. ROI-based matching 38.7% (paired mean ΔF1 = +59.7 pp [95% CI +57.6, +61.8], *d*_*z*_ = +1.67, *n* = 1,102, *p <* 0.001). At every individual condition tested, the F1 advantage was statistically significant after Holm correction (all *p <* 0.005), and S2C’s precision was uniformly *≥* 97.7% across all conditions (pooled precision *≥* 99.4% at every neuron count).

#### The ROI-based matching deficit is geometric, not footprint-driven

Giving ROI matching a perfectly stable per-neuron footprint shape did not improve its registration accuracy. Pooled across all paired runs, the best-case scenario (footprint shape fixed and unique per neuron, reused unchanged across sessions) reached ROI-matching F1 = 36.2%, essentially identical to the worst-case scenario’s 36.0%, and the S2C advantage was essentially identical between scenarios (best-case grand-pooled ΔF1 = +62.3 pp [95% CI +60.4, +64.2], *d*_*z*_ = +1.81, *n* = 1,262; worst-case = +62.4 pp [+60.5, +64.4], *d*_*z*_ = +1.81, *n* = 1,262). The two scenarios tracked each other at every neuron-count tier, with ROI F1 differing by *≤* 0.4 pp (45.9% vs. 45.6% at *N* = 100, 39.6% vs. 39.6% at 250, 33.1% vs. 33.1% at 500, and 26.3% vs. 25.9% at 1,000). Across individual (condition *×* neuron-count) cells the two scenarios differed by at most 4.3 pp with no consistent direction indicating that footprint shape jitter is downstream noise rather than a decisive identity signal. ROI matching’s failure mode on this benchmark traces to the *geometric* perturbations (rotation, translation, dropout) breaking the spatial-correlation signal upstream of any benefit footprint-shape stability could confer. The pattern is most stark at high density: at 1,000 neurons, where the mixture model collapses across the Tier-B and Tier-C stress conditions (Per-condition, the five highest-displacement conditions are B_rot Δ = +95.3 pp, B_combined Δ = +93.9 pp, B_drift Δ = +92.1 pp, C_walk Δ = +87.9 pp, and B_dropout Δ = +76.1 pp, while the three milder Tier-A conditions span Δ = +10.6 to +61.5 pp; all 32 (condition *×* neuron-count) detailed cell outputs are in Supplementary Table 1). We additionally ran both pipelines on CellReg’s publicly released sample sessions, but those sessions carry no ground-truth identity, so any comparison reduces to raw match counts that cannot be scored for correctness; we therefore confine the quantitative precision/recall comparison to the synthetic benchmark.

### 2.4 Longitudinal DMS tracking across a fentanyl behavioral economics paradigm

To evaluate the full MPS→S2C stack on a real chronic miniscope dataset, we applied the pipeline to dorsomedial striatum (DMS) calcium imaging in Long-Evans rats (*n* = 3 male, *n* = 3 female; Methods).

Raw single-photon miniscope videos were processed through MPS [18], yielding 119 *±* 27 segmented neurons per session per animal (median 111, range: 79–183; *n* = 24 sessions across 6 animals). Viral targeting of the DMS is shown in Fig. 5a. For a representative session, the same field of view is shown as a temporal mean projection after MPS preprocessing and window cropping (Fig. 5b) and the resulting MPS-extracted spatial footprint map (Fig. 5c). Event-aligned, baseline-subtracted activity traces for the most active segmented neurons in that session are shown in Fig. 5d. Lastly, S2C registered these footprints across sessions, identifying neurons recorded on different days: for a representative animal, cells tracked between two sessions (222 and 70 *µ*g/mL) are shown in green, linked across the two fields of view (Fig. 5e).

**Figure 5:**
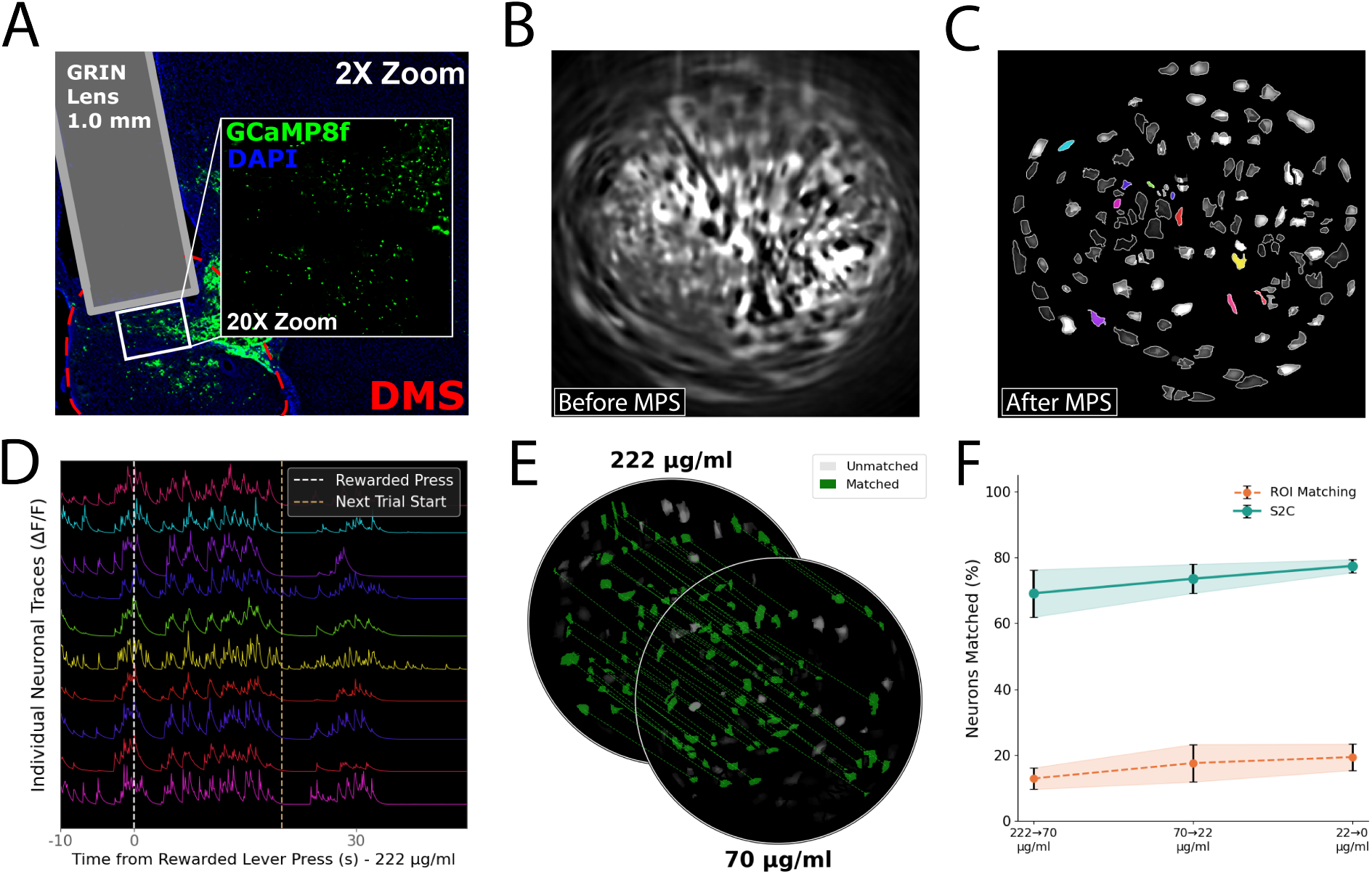
End-to-end application of S2C via same-cell longitudinal tracking of DMS neurons across an escalating fentanyl demand curve. **a**, Representative histology confirming GRIN-lens placement and GCaMP8f expression in dorsomedial striatum (DMS). Coronal section showing GCaMP8f (green) with DAPI nuclear counterstain (blue); the DMS boundary is outlined (red dashed) and the 1.0 mm GRIN-lens tract overlaid (2*×*). Inset (20*×*): high-magnification view of GCaMP8f-expressing DMS neurons beneath the lens face. **b**, Temporal mean projection of the dorsomedial striatum field of view after MPS preprocessing (deglow, denoising, and motion correction with frame removal) and window cropping. **c**, Spatial footprint map extracted from the same field of view by MPS. **d**, Baseline-subtracted activity traces for the ten most active MPS-segmented DMS neurons over a window spanning the pre-event baseline through the post-event period, aligned to a representative event (white dashed line); each trace is colored to match its footprint in **c. e**, Spatial footprint maps from two example sessions (top, 222 *µ*g/mL; bottom, 70 *µ*g/mL) for an example animal, shown as stacked, circle-cropped fields of view. Neurons matched across the two sessions are filled in green and linked by dashed lines and unmatched neurons are shown in gray. **f**, Per-step registration performance: fraction of neurons matched across each consecutive dose transition (222→70, 70→22, 22→0 *µ*g/mL) for Stars2Cells (S2C) versus ROI matching (mean *±* SEM, *n* = 6).

Across the six animals, S2C tracked DMS neurons through the four-session registration chain (222.0 →70.0 → 22.0 → 0 *µ*g/mL). At each consecutive step, S2C matched a much larger fraction of neurons than ROI matching: 69.1% *±* 7.2% vs. 13.0% *±* 3.3% (222→70), 73.6% *±* 4.4% vs. 17.6% *±* 5.7% (70→22), and 77.4% *±* 2.0% vs. 19.4% *±* 4.0% (22→0) (mean *±* SEM, *n* = 6; Fig. 5f). Because S2C registers sessions sequentially, this per-step advantage compounds: end-to-end, S2C retained 41.6% *±* 3.4% of first-session (222 *µ*g/mL) neurons across the full 222→0 chain versus 13.0% *±* 3.9% for ROI matching representing a 3.2-fold improvement in longitudinal yield. However, since GCaMP-based detection is activity-dependent, a neuron that falls silent in any intervening session cannot be segmented and drops from the sequential chain. Thus, given that per-session yield was stable across the imaging window (Methods), this 41.6% is best read as a conservative lower bound on true same-cell persistence rather than a measure of expression or indicator loss.

### 2.5 Behavioral demand and longitudinal DMS population dynamics

Animals performed oral fentanyl self-administration under a behavioral-economics demand-curve schedule (Fig. 6a, b). The resulting demand curves followed the canonical exponential decay shape, with population-level *Q*_0_ and *α* parameters consistent with published rat oral fentanyl demand work [30] (Fig. 6c; fitted *Q*_0_ = 333 *µ*g/kg, *α* = 0.00548, *R*^2^ = 0.881; P_max_ = 52.94, just beyond the tested price range). Total fentanyl intake did not differ significantly between male (*n* = 7) and female (*n* = 7) rats (Welch’s *t*(7.0) = 0.23, *p* = 0.83). Additionally, we did not observe significant changes in total locomotion across the four imaging sessions (one-way repeated-measures ANOVA, *F* (3, 15) = 0.33, *p* = 0.80, 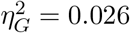, indicating the cross-session changes in demand were not driven by gross locomotor differences.

**Figure 6:**
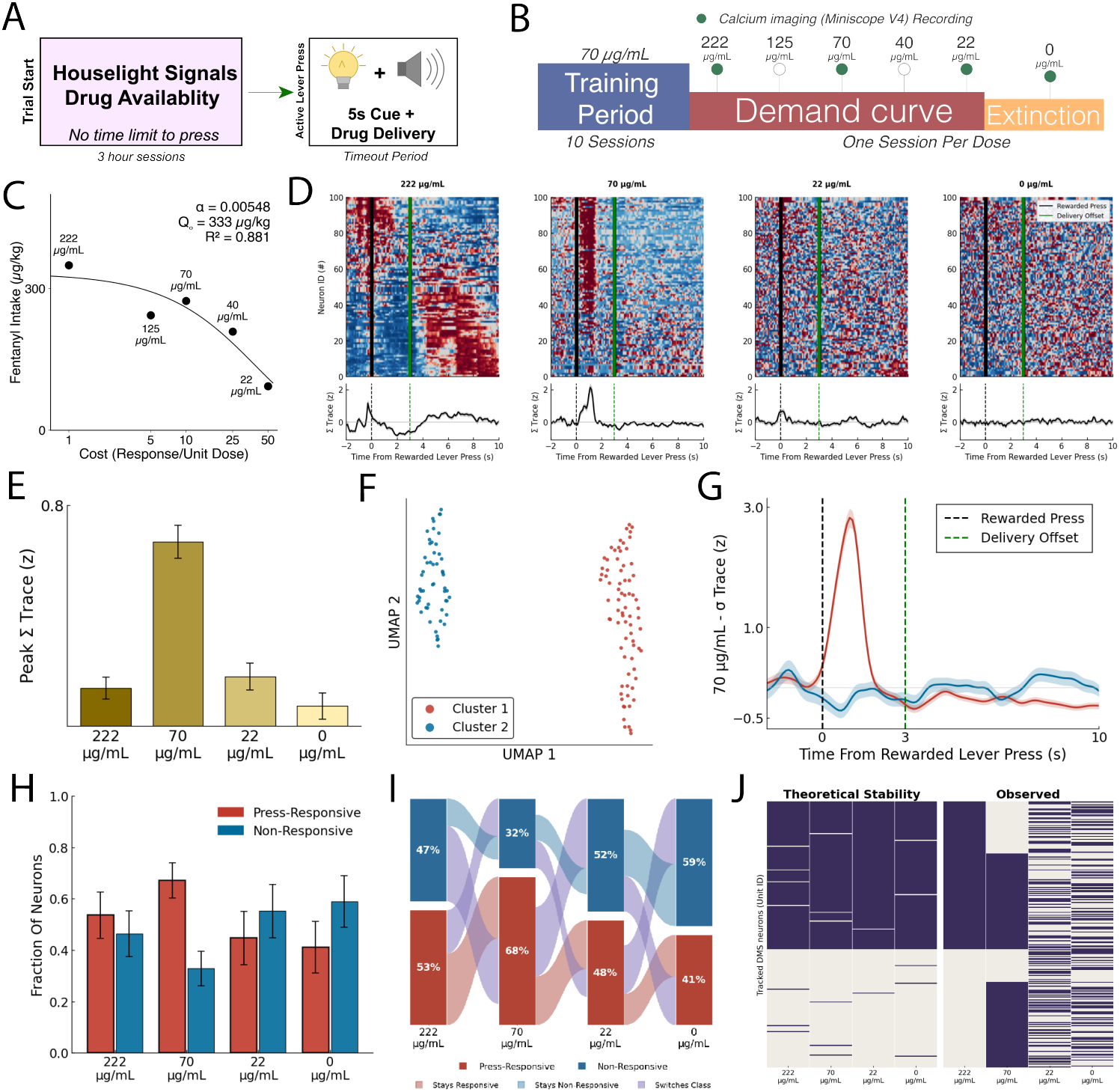
A conserved DMS population response masks near-complete single-neuron turnover across a fentanyl demand curve. **a**, Experimental timeline: habituation, acquisition at fixed concentration (70 *µ*g/mL), and across-session demand-curve schedule with descending fentanyl concentrations: 222.0, 125.0, 70.0, 40.0, 22.0 *µ*g/mL; volume held at 0.04 mL/delivery. **b**, Operant task schematic: oral fentanyl self-administration with active and inactive levers; deliveries scaled by per-session concentration to produce the unit-price gradient. **c**, Group demand curves plotted as consumption versus unit price on log-log axes; exponential demand model fit overlaid. **d**, Press-aligned DMS population activity for a representative animal at four fentanyl concentrations (222, 70, 22, 0 *µ*g/mL). For each concentration, the heatmap shows the 100 most active neurons (per-neuron *z*-score, rows ordered by temporal center of mass; early responders at top) above the population summary trace (Σ Trace; mean *z±* SEM across all segmented neurons that session); heatmaps and traces are placed on a common scale across concentrations, and dashed lines mark rewarded-press onset (*t* = 0) and delivery offset (*t* = 3 s). **e**, Group quantification of the press-aligned response across all six animals via peak Σ Trace amplitude, shaded dark-to-light by descending concentration. **f**, UMAP embedding of single-neuron press-aligned responses for the representative animal at the trained concentration (70 *µ*g/mL), each point a neuron colored by its Ward cluster. **g**, Cluster-mean response traces (mean *±*SEM, per-neuron *z*) for the two clusters in **f. h**, Fraction of segmented neurons classed press-responsive versus non-responsive at each concentration. **i**, Session-to-session flow of response-class membership for S2C-tracked neurons across the demand curve (all animals pooled). Each column is a concentration, normalized to unit height; the two bands are press-responsive and non-responsive, labeled with their fraction of tracked cells. Ribbons connect cells tracked across each consecutive pair (222 →70, 70 →22, 22 →0), colored by transition type: stays responsive, stays non-responsive, or switches class. **j**, Responsive-class identity across sessions for neurons S2C-tracked through all four imaging days. *Left (“Theoretical Stability”):* the raster the same tracking would produce if responsive identity were fixed, simulated by holding each cell’s first-session label constant and flipping it only at S2C’s per-session registration error rate (*∼*1.5%). *Right (“Observed”):* the measured labels.

To ask whether these demand differences were reflected in DMS activity, we examined rewarded lever press-aligned population responses across the concentration series (Fig. 6d). The population response was largest and most sharply defined at the trained concentration (70 *µ*g/mL), where the Σ Trace peaked at 2.13 *z* roughly 1.1 s after the press, exceeding the response at any other concentration (peak *≤* 0.66 *z*). At the 222 *µ*g/mL concentration, the press transient gave way to a brief suppression (trough *−*0.85 *z* near 2 s) and then a slow, delayed re-engagement that built after delivery offset and peaked several seconds later (0.61 *z* at *∼*6.8 s).

The response was small and tightly press-locked at 22 *µ*g/mL (0.66 *z*) and effectively absent under extinction (0 *µ*g/mL; 0.30 *z*), where the heatmap dissolved into unstructured activity. Pooling neurons across all six animals confirmed that this trained-dose enhancement was not idiosyncratic to the representative animal: the population peak response was more than threefold larger at 70 *µ*g/mL than at any other concentration (0.67 *±* 0.06 *z* vs. 0.14 *±* 0.04 at 222, 0.18 *±* 0.05 at 22, and 0.07 *±* 0.05 at 0 *µ*g/mL; mean *±* SEM, *n* = 575–777 neurons per concentration; Kruskal–Wallis *H* = 49.2, *p* = 1.2 *×* 10^*−*10^; trained vs. each, Mann–Whitney *U*, all *p ≤* 8.9 *×* 10^*−*8^; Fig. 6e).

Within each concentration, single-neuron press-aligned responses separated into two types: 1) A press-excited population and 2) A non-responsive population (Fig. 6f, g). Applying a single fixed responsiveness criterion across all concentrations, the fraction of press-responsive neurons did not differ significantly across the demand curve (Fig. 6h; 0.41–0.67 across doses; Friedman *χ*^2^(3) = 3.00, *p* = 0.39), indicating that the size of the responsive population is conserved at the group level even as price varies.

Tracking the same neurons across the demand curve revealed that the conserved responsive fraction (Fig. 6h) masked a labile underlying membership. Gains and losses between groups were balanced so the fraction held near one-half even as roughly half of tracked neurons switched between press-responsive and non-responsive at each consecutive transition (55%, 57%, and 52% for 222→70, 70→22, and 22→0; Fig. 6h, i). This turnover was of equal magnitude when concentration was held constant across consecutive 70 *µ*g/mL sessions, indicating a continuous reconfiguration of which neurons carry the press response rather than a fixed-set.

To establish that this single-neuron turnover reflects genuine reorganization rather than tracking error, we compared cross-session agreement of the responsive class against the registration ceiling the same pipeline achieves on ground-truthed data (Fig. 6j). Agreement between adjacent sessions was at chance (adjusted Rand index = +0.006; retention = 0.48 versus a 0.51 marginal responsive rate), far below the *≈* 0.94 overlap expected if responsive identity were stable. Thus, we conclude that the conserved press-responsive population (Fig. 6h) is continually rebuilt from a shifting set of neurons rather than a fixed ensemble. The turnover was of equal magnitude whether concentration rose, fell, or was held constant across consecutive 70 *µ*g/mL sessions (constant-dose ARI = +0.001), indicating ongoing session-to-session reorganization rather than a dose-locked code.

## 3 Discussion

We have presented Stars2Cells, a neuron tracking pipeline that adapts astrometric plate-solving to cross-session neuron registration in calcium imaging. By operating exclusively on centroid coordinates and using geometric descriptors invariant to rotation, translation, and uniform scaling, S2C sidesteps the requirements for spatial footprint data, activity traces, or manual alignment that limit existing methods.

### Reliable tracking plus distribution makes longitudinal analysis a default

The combination of accurate cross-session tracking and a GUI-driven distribution is what shifts longitudinal calcium-imaging analysis from a specialty technique into a standard option. Concretely, every chronic recording in an existing lab archive becomes a candidate for trajectory-level analysis: how do individual neurons reorganize their tuning across days of learning; do response profiles in reward-related circuits drift while the population code remains stable [3, 43–45]; how does ensemble identity reorganize across the time-course of a neurodegenerative phenotype; what fraction of neurons recruited on day 1 of a behavioral paradigm are still engaged on day 30. Even though none of these questions are new, for most labs, they have been mechanically out of reach. Moreover, the underlying problem of point-cloud correspondence under rigid transform with partial overlap recurs in spatial transcriptomics, developmental biology, and wildlife re-identification, and S2C is released as a general tool of which neuron tracking is the first deployed application.

### Synthetic ground truth is the only scenario that directly measures matching validation

Real-data validation of cross-session neuron tracking has a hard ceiling: there is no cell-resolved external ground truth, only signals that address fragments of the matching problem. Optogenetic tagging of a labeled subpopulation establishes positive identity for the tagged cells but provides no false-positive characterization on the unlabeled majority that comprises the bulk of any tracker’s output. Dual-channel structural markers (e.g., tdTomato co-expressed with GCaMP) supply an identity signal across sessions but are partly self-referential when the structural channel also drives the registration step [12]. Expert hand-annotation cannot integrate four-point geometric relationships across hundreds of nearly-identical neurons under arbitrary rotations and translations. The consequence is that real-data benchmarks for cross-session matching are convergent-evidence arguments, not direct measurements of matching precision and recall: they show that a tool produces consistent outputs under conditions where part of the answer is observable, but they cannot score the answer the tool actually delivers. The synthetic benchmark with construction-time ground truth is the only scenario in which matching precision and recall are directly measurable, because the identity of every neuron is known by construction and every match the tool returns can be scored as correct or incorrect. We use that approach, with parameter ranges informed by direct empirical observation of inter-session perturbations in our own and in published chronic miniscope recordings, and extended to bracket the more challenging conditions current hardware and surgical practice can produce, as the load-bearing validation step. Astrometric plate-solving in astronomy is validated identically: Astropy and Astrometry.net [19, 20] are scored against synthetically rotated star fields and ground-truthed catalogs, not against expert-by-eye matching. Thus, considering the 1,262-paired-run benchmark establishes that S2C achieves F1 = 98.4%; the DMS demand-curve application (Section 2.4) demonstrates that the pipeline carries through end-to-end on real chronic data.

### Activity cannot serve as the identity tag without encountering a tautology

Matching neurons across sessions by activity similarity assumes away the very signal longitudinal experiments exist to measure: if cells are paired because their activity is alike, one can no longer ask whether that activity changed. Drift, plasticity, learning, and addiction paradigms are designed precisely to ask how a given neuron’s activity evolves across days, and thus, geometry and not activity must carry identity (Section 2.2, Fig. 3).

### CellReg is the only like-for-like comparator

Among existing tools, CellReg is the only one that takes the same input as S2C (per-session centroids/ROIs) and targets the same task (cross-session correspondence on one-photon miniscope data), which is what makes it the natural comparator. SCOUT builds on CellReg’s spatial-correlation backbone, so CellReg upper-bounds the part of SCOUT that S2C actually competes with; SCOUT’s added temporal-correlation cue is the chance-level signal we characterize directly (Section 2.2, Fig. 3), and its remaining additions (Jensen–Shannon divergence on elliptic-Gaussian footprint fits, metric-weight perturbation, and consensus clustering) are footprint-level structural cues that require pixel footprints our centroid-only benchmark does not provide. Those are a separate, footprint-level question, not a different answer to ours. The remaining tools are not alternatives to the same problem at all: they take different inputs and answer different questions (Fig. 1c–d). Additionally, even though Centroid-only predecessors do exist STAT [46] and Coherent Point Drift [47], these methods recover identity by fitting a single spatially-smooth deformation between sessions, an assumption that degrades precisely under the miniscope recording conditions that S2C’s transform-free invariant descriptors absorb by construction. Thus, benchmarking any of these orthogonal methods against S2C would score them on a task they were never built for, so we situate them in the layer diagram rather than treat them as head-to-head competitors.

### ROI matching fails on geometry, not footprint quality

Even an idealized ROI matcher with perfectly stable per-neuron footprints cannot recover from the inter-session FOV variability that chronic single-photon recordings routinely produce. Thus, this is precisely the variability a rotation-invariant geometric approach is built to absorb, which is why holding footprint shape fixed (best-vs. worst-case) barely changes ROI accuracy (Fig. 4g).

### S2C is purpose-built for the chronic-imaging density approach

We tested S2C across neuron-count tiers from 100 to 1,000, which spans the standard density range for chronic miniscope recordings (typically 50–200 neurons per FOV in deep-brain GRIN-lens preparations) and standard two-photon cortical imaging (typically 200–1,000 neurons per FOV). High-end mesoscope-class two-photon recordings, which can reach 3,000–12,000+ neurons per FOV [48–50] – are outside this scope by design. Cross-session tracking at that density is a different engineering problem with different bottlenecks: descriptor count grows as *O*(*N* ^2^), the 4D descriptor space saturates, and the false-positive descriptor matching rate at fixed 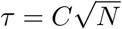 rises. Adapting S2C to that approach is a tractable extension (locality-sensitive hashing in descriptor space, hierarchical RANSAC, GPU batching of inlier evaluation) rather than a new method. We have not attempted that extension here because the dominant unsolved adoption gap sits at densities S2C already covers.

### From benchmark to biology

Beyond the synthetic benchmark, the DMS fentanyl behavioral-economics application (Section 2.4) demonstrates that S2C recovers biologically meaningful structure from real chronic data. ROI-based matching recovered only 13.0% *±* 3.9% of first-session neurons across the full 222→0 *µ*g/mL chain versus 41.6% *±* 3.4% for S2C, a 3.2-fold difference. Thus, roughly two-thirds of the longitudinally trackable population, and whatever across-session dynamics those cells carry, would have been discarded as untrackable via solely ROI matching.

### A stable DMS population code conceals single-cell turnover

Within each dose, DMS neurons split into a press-responsive group and a non-responsive remainder, classed by the response shape (Fig. 6f, g) staying essentially constant across the demand curve experiment (Fig. 6h; Friedman *χ*^2^(3) = 3.00, *p* = 0.39). Notably, the same cells do not stay in their group across sessions. Instead, neurons drop out of and rejoin the responsive group between consecutive doses (Fig. 6i) shuffling at such a rate that a cell’s group at one dose predicts its type at the next no better than chance (adjusted Rand index *≈* 0). It is important to note that since: I) A stable identity would yield *≈* 0.94 agreement through the same pipeline, far above the observed labels (Fig. 6j) and II) The turnover both persists at constant dose (70 *µ*g/mL) and survives a noise-robust graded-response measure, we can rule out a binary-threshold artifact. Thus, the observed conserved press-responsive population does not represent a fixed ensemble but one continually rebuilt from a shifting set of neurons where the group-level signal holds steady while its single-cell membership turns over completely. This dissociation is the signature of representational drift, established in hippocampus, cortex, and piriform and now extended to the striatum [3, 5, 35, 43, 44]. Pairing this same-cell tracking with D1/D2 reporters would resolve whether the cells cycling through the responsive group do so within or across striatal pathways, a natural next step beyond this proof-of-concept. Nonetheless, since a bulk signal sums across cells, this turnover would have been invisible at the aggregate level – requiring single-cell imaging with same-cell tracking. Moreover, geometry-carried identity is what made this distinction measurable since activity-based matching would assume the identity it exists to test and ROI matching misses the pairs outright.

### Limitations

S2C’s accuracy depends on centroid extraction quality from the upstream pipeline; systematic localization errors will propagate. The current implementation assumes rigid (or similarity) inter-session transforms; large nonlinear distortions are handled only to the extent they appear locally rigid within quad neighborhoods. The descriptor pipeline is *O*(*NkK*^2^) (linear in *N* at fixed *k* and *K* ) and descriptor matching is *O*(*N* ^2^), the latter the binding cost at high density. Regardless, within the tested ranges, S2C did not drop below 93.6% F1 on any condition, and we therefore cannot point to an empirically observed sub-90% breaking point. Two failure modes are predictable from the algorithm’s construction, however, and worth flagging for users whose preparations sit outside the chronic-rodent-imaging scope this paper addresses. First, when nearest-neighbor distance approaches the centroid jitter floor, then quads from different neighborhoods become descriptor-similar and the matching step’s signal-to-noise floor could break down. Second, because our RANSAC step fits rigid transforms (rotation + translation) with scale held fixed, it rejects genuine matches in preparations where tissue expands uniformly between sessions, such as developmental two-photon recordings, organoids, and plant-growth tracking. However, releasing the scale parameter inside RANSAC is a single-line code change, but we have not implemented or validated it since it is outside the scope of the present paper. Future work could also incorporate locally affine models or deep-learned descriptors to handle nonlinear distortions more explicitly; the higher-density extension is outlined above.

### Closing

Thousands of investigators now image neurons in behaving animals, but only a small minority can follow the same cells across days. Stars2Cells adds that capability on top of the segmentation pipelines these recordings already rely on. Thus, once individual neurons are linked across sessions, a chronic recording stops being a set of isolated snapshots and becomes a record of how the same cells change over an untold number of modalities.

## 4 Methods

### 4.1 Stars2Cells algorithm

S2C solves the same problem astronomers solve when plate-solving a photograph of an unknown patch of sky: given a sparse set of points in unknown rotation and translation, with some points missing and some new ones appearing, identify which point is which. The solution is borrowed from astrometric quad matching [20] and lent to neurons in the name S2C in that no individual point carries enough information to be identified, but four points carry much more. The relative geometry of any four neurons is invariant under rotation and translation, and once that geometry is encoded as a single vector in ℝ^4^, two sessions’ quads can be compared directly without ever solving for the unknown transform. Once enough quads agree, the transform falls out of them.

S2C was implemented in Python 3.10 against a pinned dependency stack: NumPy 1.26.4 and SciPy 1.13.0 (numerical core, installed via Conda), faiss-cpu (cosine-distance descriptor matching), scikit-learn 1.8, scikit-image 0.26, NetworkX 3.6, tqdm 4.67, psutil 7.0, and PyQt5 5.15 / PyQt5-sip 12.18 / pyqtgraph 0.14 for the GUI; image and array I/O via Pillow 12.1, ImageIO 2.37, tifffile 2026.3.3, and Matplotlib 3.10. The full pinned requirements.txt ships with the source release. Both builds are produced with PyInstaller, which bundles a self-contained Python runtime and all scientific dependencies into the application itself: the macOS build is delivered as a signed and notarized DMG containing Stars2Cells.app, and the Windows build is delivered as a ZIP archive containing a standalone Stars2Cells.exe. End users do not need to install Python, manage Conda or virtual environments, edit PATH, or use the command line at any point in the workflow. Standard operating-system gatekeeping warnings (macOS Gatekeeper, Windows SmartScreen) may appear on first launch and are dismissed through the OS’s standard install-confirmation flow, requiring no command-line intervention.

#### Step 1: Quad descriptor generation

Quads are not enumerated brute-force 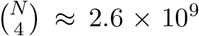 candidates at *N* = 500 is intractable, and almost all of them are uninformative (very thin or very small quads carry essentially no positional information). S2C builds quads diagonal-first and draws each quad exactly once.

The diagonal set per session is constructed from *k*_local_ = min(15, *N −* 1) (tunable) KD-tree nearest neighbors plus *k*_random_ = max(2, *k*_local_*/*2) random long-range partners (tunable) per neuron (with KNN exclusions, seeded for reproducibility). Local diagonals stabilize the descriptor under small inter-session shifts; long-range diagonals produce FOV-spanning quads with small descriptor blur (*σ*_*d*_ *∝* 1*/L*_*q*_, where *L*_*q*_ is the longest pairwise distance of the quad) and dominate matching when the imaging field shifts substantially between sessions. For each unique diagonal (*i, j*) with *i < j*, the signed perpendicular height of every other neuron *k* is computed inline via the 2D cross product:

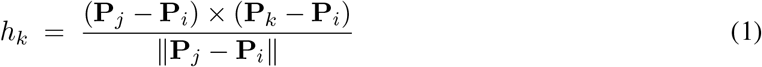

where **P**_*i*_ *∈* ℝ^2^ is the centroid coordinate of neuron *i* and *×* denotes the scalar 2D cross product (**u** *×* **v** = *u*_*x*_*v*_*y*_ *−u*_*y*_*v*_*x*_). The top-*K* third points by |*h*_*k*_| are retained (*K* = 25 default) and combined into 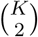 candidate quads per diagonal. For each quad *{i, j, C, D}*, the four-dimensional descriptor is constructed by projecting *C* and *D* onto the diagonal-aligned coordinate frame:

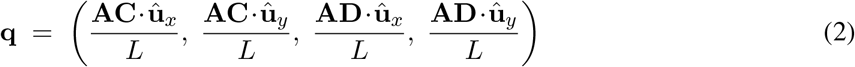

where **A, B** are the two endpoints of the quad’s longest pairwise edge, *L* = *∥***B** *−* **A***∥*, **û**_*x*_ = (**B** *−* **A**)*/L*, and **û**_*y*_ is its 90^*•*^ counter-clockwise rotation. Canonical ordering enforces *x*_*C*_ *≤ x*_*D*_ (swap labels otherwise) and (*x*_*C*_ + *x*_*D*_) *≤* 1 (axis flip otherwise); these two rules are what convert the raw projection from rotation-equivariant to fully rotation-invariant. Degenerate quads with |*y*_*C*_|, |*y*_*D*_| *<* 10^*−*4^ (collinear) are discarded.

The same quad has six pairwise edges and could in principle be reached from any of them. The **ownership guard** emits a quad from a diagonal only when that diagonal equals the quad’s longest edge, evaluated within a 10^*−*6^ relative tolerance. This yields exactly one emission per quad with no global deduplication pass. For real-valued centroid coordinates, two distinct pairwise distances are equal only on a measure-zero set, so each quad has a unique longest edge almost surely; the 10^*−*6^ relative tolerance in the ownership test collapses the rare floating-point near-tie to a single emission.

### Coverage remediation

The ownership guard can miss a neuron in a dense region where every quad incident to it gets claimed by some longer neighboring diagonal, and thus, the neuron contributes zero descriptors. To solve this, after the main pass, S2C identifies neurons whose quad count falls below a tunable parameter (default = 0.4) multiplied by the median(coverage) over the field and re-runs descriptor generation on their incident diagonals with the ownership guard disabled. The remediation batch is deduplicated against the existing quad set on sorted vertex tuples and quality-pruned by |y_*C*_| + |*y*_*D*_| (keeping the top fraction, default 1.0 – tunable). Formal detection criteria and the uniqueness handling for the remediation batch are given in Supplementary Note 3.

#### Descriptor blur and saturation

Per-neuron centroid jitter of standard deviation *σ*_*j*_ pixels propagates to descriptor space as 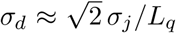, where *L*_*q*_ is the quad’s diagonal length: each projected descriptor coordinate involves a difference of two jittered points (the 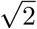factor) normalized by *L*_*q*_. Longer-range diagonals therefore produce smaller descriptor blur, which is why the diagonal set mixes KNN-local edges with random long-range partners. The descriptor field is *saturated* when median nearest-neighbor distance in descriptor space falls below median descriptor blur, median(NN_*d*_) *<* median(*σ*_*d*_); in that scenario jitter-induced descriptor noise exceeds inter-descriptor separation and false matches are unavoidable regardless of threshold. S2C evaluates this criterion at the end of Step 1 and emits a warning if saturation is detected.

#### Step 1.5: Threshold calibration

Whether two quads count as the same comes down to a distance threshold in ℝ^4^ descriptor space: tight enough to suppress chance collisions between unrelated quads, but loose enough to absorb the centroid jitter that survives MPS preprocessing. Both moving parts depend on *N* : as the field densifies, the local-diagonal lengths that anchor same-cell descriptors shrink, so the same centroid jitter occupies a larger fraction of each diagonal and same-cell descriptor blur grows. The threshold that must clear that blur therefore grows with field size.

For each animal, S2C samples a fixed number of session pairs (default 10k quads per pair – tunable) and sweeps similarity thresholds *τ ∈* [0, 1] using descriptor distance only. At each *τ*, raw descriptor matches are passed through a consistency filter, and a quality score is computed from the raw and filtered counts. The optimum *τ*^*\**^ is selected as the location of the per-pair quality peak. The relationship

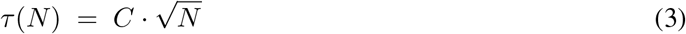

is then fit by least squares across all sampled pairs from that animal, yielding *C, C*_std_, and *R*^2^. The 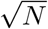form follows from a geometric argument (Supplementary Note 2): local-diagonal length scales as *L*_*q*_ *∝ N*^*−*1*/*2^, so same-cell descriptor blur scales as 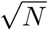, and *τ* is bounded below by it. Because the derivation assumes uniform density, the regression *R*^2^ reports how closely a field approximates uniform packing, not calibration quality: a lower *R*^2^ simply means centroids are non-uniformly spaced (clustering or density gradients, both expected), with the residual imprecision absorbed downstream by the consistency filter and RANSAC.

#### Step 2: Descriptor matching

For each session pair (*R, T* ), every quad descriptor in *T* is matched to its top-1 nearest neighbor in *R* under cosine distance via FAISS (IndexFlatIP with L2-normalized descriptors), and matches with distance below the per-animal calibrated *τ* are retained. A geometric consistency filter then rejects matches whose ratio of inter-quad centroid distances is inconsistent with the bulk distribution (consistency threshold = 0.8 – tunable). Surviving matches are written to a lightweight NPZ file containing the matched quad indices and the underlying centroid coordinates.

#### Step 2.5: RANSAC geometric verification

The output of Step 2 contains real correspondences and chance-collision noise in unknown ratio. The structure that distinguishes them: every real match is consistent with a single rigid transformation (**A, t**) between the two sessions, and every fake match is not. S2C draws *K* = 1,000 (tunable) random subsets of *m* = 4 matched centroid pairs each, and estimates all *K* rigid transforms simultaneously via batched SVD on the cross-covariance matrices:

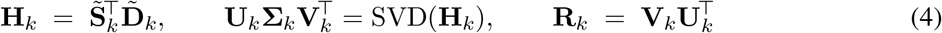

where 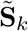 and 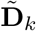 are the centered source and destination samples of the *k*-th subset. Reflections are corrected by flipping the last column of **V**_*k*_ whenever det **R**_*k*_ *<* 0, after which the rotation is recomputed. Translation is recovered as 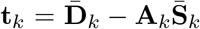 where the bars denote sample means. Inlier counts for all *K* candidate transforms are evaluated against all *N* matched pairs in float32 chunks sized to *∼* 200 MB of working memory, with squared-residual threshold (5.0 px - tunable)^2^. The transform with the most inliers is then re-estimated on its full inlier set, and matches with refined residual *>* 5.0 px (tunable) are rejected. The RANSAC residual threshold used is persisted into the Step 2.5 NPZ output, where it is consumed by Step 3 to set its distance cutoffs.

#### Step 3: Hungarian assignment and consolidation

Step 2.5 leaves a set of *quad*-level correspondences that are geometrically consistent with the inferred rigid transform. Step 3 turns these into a one-to-one *neuron* mapping. Each surviving inlier quad votes for all 4 *×* 4 = 16 implied neuron pairings, building a vote matrix 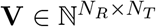 . Raw vote counts are degree-normalized to remove a hub-neuron bias:

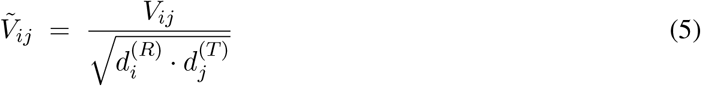

where 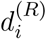 and 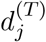 are the total quad degrees of neurons *i* and *j* in their respective sessions. The cost matrix combines vote-cost and spatial residual under the inferred transform:

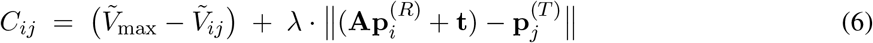

with *λ* chosen so that the median spatial term equals the median vote term over voted entries 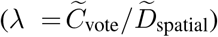, and *C*_*ij*_ = *∞* outside a distance cutoff of 3.0*×* (tunable) the RANSAC residual threshold. Pairs that are spatially nearby but unvoted are not blocked outright; they receive a distance-only fallback cost above the maximum voted cost, so they remain assignable when no voted competitor exists.

The cost matrix is then padded with rectangular dummy rows and columns (one dummy slot per real neuron in each direction, each priced at the global median voted cost) and solved with SciPy’s linear sum assignment routine (Pass 1). Without the dummies the Hungarian algorithm would return a complete matching that pairs every neuron to some partner; the fixed-cost dummy slots instead give each neuron an opt-out, so it accepts a real partner only when that partner is cheaper than the median dummy cost and is otherwise left unmatched, which is the correct behavior for dropouts and newly appearing cells that have no true correspondent. Pass-1 matches with post-hoc spatial residual exceeding 1.0*×* the RANSAC threshold are rejected. Pass 2 then re-solves a smaller Hungarian over the unmatched residual 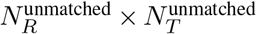 submatrix) under a relaxed cutoff of 2.0*×* the RANSAC threshold, with submatrix dummy cost set to the 75^th^ percentile of finite submatrix costs (tunable). This recovers correspondences that Pass 1 conservatively rejected.

Each surviving match (*i, j*) is assigned a confidence score in [0, 1] that averages four equally-weighted components: the degree-normalized vote score 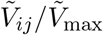; spatial proximity max(0, 1 *− d*_*ij*_*/d*_cutoff_); assignment margin 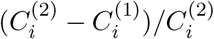 where 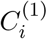 and 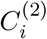 are the best and second-best finite costs in row *i*; and pass score (1.0 for Pass 1, 0.5 for Pass 2). Pairwise neuron correspondences are then chained transitively into multi-session tracks: a track threading neurons *n*_1_ *→ n*_2_ *→ · · · → n*_*S*_ across *S* sessions inherits the minimum link confidence as its conservative track confidence, and the mean as its expected confidence.

### 4.2 Temporal correlation control experiment

To test whether calcium-trace correlation could serve as a stand-alone identity signal, we generated 1,000 simulated neurons placed at identical spatial positions (Poisson-disk, *r*_min_ as in the main benchmark) across 5 sessions. For each session, calcium traces were independently simulated using an AR(1) model: perneuron Poisson spike trains with rates *∼ U* (0.05, 0.5) Hz at 10 Hz frame rate were convolved with a single-exponential calcium kernel (*τ* = 1 s) using a discrete IIR filter with coefficient *a* = *e*^*−*1*/*(*τ·*fps)^, scaled by per-neuron amplitudes *∼ U* (0.5, 3.0), and corrupted with Gaussian noise (*σ* = 0.05). The random seed for trace generation differed between sessions, so the dynamics of any given neuron were statistically independent across sessions even though its position was perfectly preserved. Recording durations of 10, 20, 60, and 180 min were tested (6,000 / 12,000 / 36,000 / 108,000 frames). For each pair of sessions, traces were *z*-scored row-wise (in float32) and the full 1,000 *×* 1,000 Pearson correlation matrix was computed in row-chunks of 100 to avoid the *∼*8 GB peak of a single dense matmul; each neuron in session *s*_1_ was assigned to its maximally correlated counterpart in session *s*_2_. Accuracy was scored against the (preserved) ground-truth identity. Across all 10 session pairs and all four durations, mean accuracy was indistinguishable from chance (1*/N* = 0.1%).

### 4.3 Synthetic benchmark generation

Benchmark data were generated by a custom Python script that placed neurons via Bridson Poisson-disk sampling and applied deterministic affine transforms with NumPy under controlled per-condition seeds. For each combination of *animal* (8), *condition* (1–9 depending on tier; see below), *seed* (5; except Tier 0, single seed), and *neuron count* (100, 250, 500, 1,000), five consecutive sessions were generated from a base centroid field placed in a 600 *×* 600-pixel FOV with *r*_min_ set to 55% of the theoretical hex-packing maximum.

#### Per-session transformation pipeline

Each session is generated from the base field by a fixed transform sequence detailed in Supplementary Note 5 (cumulative rotation, translation, clip-to-FOV, dropout, per-neuron jitter, ID permutation), with ground-truth identity captured before permutation.

#### Neuron-count tier coverage

All four neuron counts (100, 250, 500, 1,000) run all four tiers (Tier 0 + A + B + C; 8 productive conditions plus 1 identity sanity-floor).

### 4.4 Spatial footprint synthesis for the ROI-based matching comparison

ROI-based matching requires pixel-level spatial footprints (ROI masks), not just centroids. To enable a like-for-like comparison without a lossy round-trip through real movies, we synthesized footprints directly from the same centroid fields used to evaluate S2C. Each neuron was rendered as a 2D elliptical Gaussian with semi-major axis *σ*_major_, semi-minor axis *σ*_minor_ = *σ*_major_ *· ϵ* (eccentricity *ϵ ∼ U* (0.6, 1.0)), orientation *θ ∼ U* (0, *π*), and peak intensity *∼ U* (0.6, 1.0). The mean radius was scaled with neuron count to mimic empirically observed CNMF/CNMF-E footprint compression in dense fields: 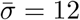 px at *N* = 100, 10 px at 250, 8 px at 500, and 6 px at 1,000 – values consistent with the *∼*8–15 px effective radii reported for one-photon miniscope CNMF-E extraction [8, 51] and the FOV-density scaling used by CNMF-E itself. Per-neuron radius variability was CV = 0.15, clipped to 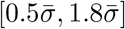. Footprints were rendered with a 2.5*σ* crop radius for efficiency and stored both as compact per-neuron patches (with bounding-box offsets) and as dense *N × H × W* arrays consumable directly by CellReg in either Python (.npy) or MATLAB (.mat, with *H × W × N* axis order) formats. Each footprint was L2-normalized.

#### Two scenarios

Footprints were generated under two scenarios designed to bracket ROI matching’s plausible operating range:

- **Best-case (no shape perturbation)**. The footprint shape parameters of each neuron are fixed once per animal, are unique per neuron, and reused unchanged across every session, condition, and seed; only centroid *position* changes between sessions according to the benchmark’s geometric perturbations. This gives ROI matching the strongest possible spatial-correlation signal: a neuron’s footprint shape is a perfect identity tag.
- **Worst-case (complete shape perturbation)**. Per-session, per-neuron Gaussian perturbations are added on top of the base shape parameters: *σ* jitter *∼ N* (0, 0.5 px) (independent on major and minor axes), orientation jitter *∼ N* (0, 5°), peak-intensity jitter *∼ N* (0, 5%). These magnitudes were chosen to model focus drift, motion-correction residuals, CNMF re-estimation noise, and slow expression/photobleaching changes that real footprints undergo across days.

The same per-animal base-shape parameters are reused across all S2C-vs-ROI-matching comparisons within each scenario, so the spatial signal that the ROI-based matching sees is held constant within a scenario and varies only through the explicitly imposed per-session perturbations.

### 4.5 ROI-based matching reimplementation

To enable evaluation across all 32 productive (condition *×* neuron-count) cells on a single workstation, we reimplemented CellReg’s core registration pipeline in Python (NumPy, SciPy). The reimplementation faithfully ports the pipeline component-for-component, in the order Sheintuch et al. specify it:

1. **Footprint normalization and FOV alignment**. Each footprint is L2-normalized to unit sum (matching MATLAB’s normalize_spatial_footprints.m), and sessions are zero-padded to a common canvas. A pre-matching alignment stage (align_images) estimates the rigid-body transform that maximizes cross-correlation between each non-reference session’s footprint projection and the reference, then applies that transform to both footprints and centroids. We retained CellReg’s full alignment step so that the comparison is not unfairly handicapped by inter-session shifts the tool is designed to absorb upstream of matching.
2. **All-to-all neighbor distributions**. For each cell pair across sessions within a maximal centroid distance *d*_max_ (default 20 *µ*m at 1 *µ*m/px, so 20 px), the spatial Pearson correlation *ρ*_spatial_ is computed over the union of non-zero pixels (corr2-equivalent), and the centroid distance *d* is recorded. The pooled (*ρ, d*) distributions across all session pairs are accumulated for model fitting.
3. **Two parallel mixture models on** (1 *− ρ*) **and on** *d*. A spatial-correlation model fits a *lognormal* same-cell distribution and a *Beta* different-cell distribution to the (1 *− ρ*_spatial_) histogram via 100 EM iterations, with sigmoid smoothing on the same-cell component (matching MATLAB CellReg). A centroid-distance model fits a lognormal same-cell + sigmoid-by-linear different-cell mixture to *d* via nonlinear least squares. Each model produces a Bayes-rule lookup *P* (same) as a function of its metric, and the intersection (where *P* (same) = 0.5) is reported.
4. **Best-model selection**. choose_best_model picks the model with the lower combined cost cost = FP_rate + FN_rate + MSE.
5. **Per-pair** *P*_**same**_ **assignment**. Each candidate neighbor pair is given its *P*_same_ via nearest-bin lookup in the chosen model.
6. **Initial registration – single greedy pass**. The registry is seeded with all cells from session 1; for sessions 2, …, *S*, each cell is greedily matched to its highest-similarity registered neighbor above the threshold (or registered as a new cell otherwise). Slot conflicts are resolved by keeping the higher-scoring assignment and displacing the loser to a new row. *This is not a Hungarian solve*: it is a one-pass nearest-neighbor seed (initial_registration_*).
7. **Final registration – iterative cluster_cells (***≤***10 iterations)**. The greedy seed is refined by the iterative cluster_cells routine. Each iteration performs three sub-steps: (A) per-cell reassignment within 1.7 *× d*_max_ using the maximal-similarity criterion (split clusters whose best similarity falls below threshold; move or swap cells between clusters when a better-fitting cluster is found); (B) cluster merging across non-overlapping session sets when the maximal pairwise *P*_same_ exceeds threshold; (C) cleanup of empty rows. The loop terminates when fewer than 10 reassignments occur in an iteration or after 10 iterations.
8. **Cell-level scoring**. Each registered cluster receives positive (mean *P*_same_ over confirmed pairs), negative (mean 1 *− P*_same_ over absent-session pairs), and exclusive (penalized for spurious extra neighbors) scores via compute_scores.

#### Configuration parameters

Microns per pixel = 1.0; maximal pairing distance = 20 *µ*m; alignment = “Translations + Rotations”; *P*_same_ threshold = 0.5. These settings match standard CellReg defaults for miniscope data.

#### Search-radius choice

The 20 *µ*m radius defines the neighborhood within which CellReg’s two mixtures are fitted: the same-cell mode (lognormal on centroid distance; lognormal on 1 *− ρ*_spatial_) sits at small values and the different-cell mode is fitted across the remainder of the radius, with the discriminative power of *P*_same_ deriving from the gap between them. Extending the radius to admit additional candidate matches in the highest-displacement conditions would flood the neighbor pool with different-cell pairs at every distance, compress the gap between the two modes, and produce low-precision matches under an identity model that has ceased to mean what CellReg’s authors specified. We therefore retained the standard miniscope-default radius across the full benchmark. When per-step inter-session displacement exceeded that radius, the same-cell mode of CellReg’s centroid-distance mixture lost within-radius support and only chance pairings between unrelated cells remained (TP = 0, F1 undefined); these runs were excluded from the paired statistic rather than recovered by parameter widening, and they concentrate in the B-drift condition, whose 5–25 px/step translation range crosses the 20 *µ*m horizon directly. Because these excluded runs are precisely the cases where ROI-based matching failed completely, the exclusion is conservative with respect to our central comparison: had we instead scored them as F1 = 0, CellReg’s pooled F1 would fall and the S2C advantage would widen. Dropping them therefore understates rather than inflates the reported gap.

To verify the reimplementation, we generated held-out 2-session test sets at all four benchmark neuron counts (*N* = 100, 250, 500, 1,000) using generate_cellreg_test.py, each with *∼*1 px per-neuron centroid jitter and 10% session-2 dropout, with permuted ROI IDs and a separately stored ground-truth identity table mapping each session’s permuted indices back to base identity. Identical synthetic spatial footprints were exported in both formats: .npy dictionaries for the Python implementation and .mat arrays in *H × W × N* orientation for native MATLAB CellReg (v1.5.9, which incorporates the FOV-alignment fix introduced in v1.5.5). Both pipelines were run end-to-end with matched parameters (microns/pixel = 1.0, maximal distance = 20 *µ*m, *P*_same_ = 0.5, “Translations + Rotations” alignment, “Spatial correlation” model selection at both intermediate and final steps), and the resulting cell-to-index maps were compared row-by-row against ground truth.

At every neuron count tested (*N* = 100, 250, 500, 1,000), the two implementations agreed on *every* matched pair: identical model selection (spatial correlation), identical match count, identical pair assignments, identical singleton (session-1-only) classification of the dropped neurons, and identical 100% precision and 100% recall against ground truth. The set difference between Python and MATLAB matched-pair sets was empty in both directions across all four neuron counts. With pair-level concordance established across the full neuron-count range of the benchmark, the parallelized Python reimplementation was used for the full benchmark. Because the implementation is deterministic and its code path does not branch on input difficulty, the pair-level concordance established on these benign test sets carries over to the harder benchmark conditions; we reimplemented in Python solely because running native MATLAB CellReg across all 32 (condition *×* neuron-count) cells at scale on a single workstation was infeasible, not to alter its behavior.

#### Performance under benign conditions

Pair-for-pair agreement with native MATLAB CellReg is established in Methods (“ROI-based matching reimplementation”). A one-to-one reproduction of their reported numbers is not attainable: Sheintuch et al. report false-positive/false-negative rates rather than F1, the only public CellReg data are unlabeled (precluding scored precision/recall), and their real-recording error rate cannot be matched by clean synthetic data without tuning density and noise to the target, which is circular. We therefore rely on the agreement with the released MATLAB tool as the rigorous check. As a qualitative consistency test, on well-aligned synthetic data the reimplementation registers cells with *<*5% error, consistent with the *<*5% rates Sheintuch et al. report [9].

### 4.6 Evaluation metrics

Precision, recall, and F1 were computed from ground-truth identity labels. A match was scored as true positive if both neurons shared the same base identity (accounting for permutation). For multi-session chains, metrics were computed pairwise across all four consecutive transitions (A→B, B→C, C→D, D→E). We selected F1 as the headline metric since both error modes carry biological cost in this setting: a false positive corrupts a neuron’s apparent dynamics by binding it to traces from a different cell, while a false negative discards a real cross-session correspondence and reduces the longitudinal sample. F1 weights both equally, which we considered preferable to weighted variants (e.g., F_*β*_) that would require choosing *β* on grounds we have no principled basis to set.

### 4.7 Bibliometric adoption analysis

To quantify the longitudinal-tracking adoption gap reported in Fig. 1 and the Introduction, we performed a programmatic bibliometric survey using the NCBI Entrez API (BioPython 1.83) for publication counts and the OpenAlex API for tool-citation counts, on a date locked to manuscript submission. For the PubMed queries, all matching PMIDs were retrieved via esearch, full record summaries were pulled via esummary in 500-ID batches, and unique Last Author fields were tallied. Three measures were taken: (i) “two-photon labs” (all years, ẗwo-photon[̈Title/Abstract]), yielding 8,484 unique last authors from 19,642 unique PMIDs (full uncapped retrieval); (ii) “longitudinal labs” (2021+, l̈ongitudinal[̈Title/Abstract] AND any of c̈alcium imaging̈, ẗwo-photon̈, m̈iniscope,̈ öne-photon̈, ẗracking neurons,̈ c̈ell registrationïn Title/Abstract); and (iii) tool-specific adoption, estimated as citations of each tool’s canonical reference publication via the OpenAlex API, using manually verified DOIs. We report citation counts because no single index captures software adoption completely: PubMed title/abstract queries miss tools cited only in Methods (PubMed indexes no full text); open-access full-text indices (Europe PMC) recover Methods/Results mentions but only for the open-access subset; and citation counts, while covering the full literature irrespective of access status, conflate citation with use and under-count tools distributed without a citable paper. Counts are therefore transparent order-of-magnitude estimates; tools published within approximately one year of access (CaliAli, Track2p) are correspondingly under-represented and reflect recency rather than limited utility. Active-laboratory population estimates for UCLA Miniscope (*∼*500 labs) and Inscopix nVista/nVoke systems (*∼*600 labs) are taken as *manufacturer/ community estimates as of April 2026*, sourced from the UCLA Miniscope BRAIN Initiative ToolMakers page [52] and reporting on the Bruker–Inscopix acquisition [53], respectively.

### 4.8 Animals and housing

For all experiments, 8-week-old male and female Long-Evans rats were bred in-house. Rats were dual-housed in a temperature- and humidity-controlled room on a 10:14 dark-light cycle and acclimated to the facility for *∼*1 week before beginning experimental procedures at 9–10 weeks old. Food and water were available ad libitum throughout the experiment (weight range: 250–450 g). Experimental procedures were approved by the Institutional Animal Care and Use Committee for the Veteran’s Affairs, Puget Sound Health Care System

#### ARRIVE 2.0 reporting elements

A total of *n* = 6 animals (3 male, 3 female) were used in the imaging cohort; this sample size was chosen to match the per-cohort *n* used in the precedent oral fentanyl behavioral-economics work in the same lab [30] and is treated here as a proof-of-concept demonstration of the MPS→S2C stack on real chronic miniscope data rather than as a powered biological-discovery experiment.

Behavioral sessions were run in fixed-time daily blocks; imaging-day order was fixed by the demand-curve schedule (sessions 12, 14, 16, and 17). *Exclusion criteria, pre-specified:* animals were excluded from analysis if (i) viral expression was outside the dorsomedial striatum on histology, (ii) the GRIN-lens placement was outside the dorsomedial striatum on histology, (iii) baseline operant responding failed to stabilize during acquisition, or (iv) the miniscope FOV was unusable (lens occlusion, severe motion artifact uncorrectable by MPS, or signal loss).

### 4.9 Oral fentanyl self-administration and behavioral economics

Animals (n = 7 male, n = 7 female) were trained on an oral fentanyl self-administration paradigm following the procedure of [30], with concentrations and volumes adjusted for the behavioral-economics demand-curve schedule used here; of these, a subset of 6 (n = 3 male, n = 3 female) were implanted with a GRIN lens and miniscope baseplate and constitute the imaging cohort analyzed throughout (see “Surgical preparation and chronic miniscope imaging”). Training was conducted at a fixed concentration of 70 *µ*g/mL, with each delivery providing 0.04 mL of solution. After acquisition and stable responding on the training concentration (sessions 5–9: 56.0 *±* 9.2 active-lever presses per session, mean *±* SEM, *n* = 15; Friedman *χ*^2^(4) = 2.79, *p* = 0.59), animals advanced to an across-session demand-curve schedule, in which the fentanyl concentration descended across days while the response cost per unit dose increased monotonically. The per-session concentrations (in *µ*g/mL) were 222.0, 125.0, 70.0, 40.0, and 22.0, followed by extinction at 0 *µ*g/mL, with delivery volume held constant at 0.04 mL.

For each delivery, the dose in micrograms per kilogram was calculated as the per-session fentanyl concentration multiplied by the per-delivery volume (0.04 mL) and divided by the animal’s body weight in kilograms. Whole-brain fentanyl concentration over time was estimated using the same two-compartment model previously applied to oral fentanyl in rat [30], adapted from the cocaine pharmacokinetics methodology of [54] with a larger elimination coefficient (*k*_el_ = 0.175) to account for fentanyl’s relatively longer half-life.

Briefly, each delivery contributed a bi-exponential term with a fast distribution component and a slower elimination component, parameterized by a transfer constant *k*, an apparent brain volume *v*, and rate constants *α* and *β*. In practice, this was implemented as a convolution of the delivery times with a kernel proportional to exp(*−α*Δ*t*) *−* exp(*−β*Δ*t*), where Δ*t* is the time since each prior delivery. Concentrations were evaluated on a dense temporal grid spanning the full 180-minute session for each animal and then averaged across animals to obtain a population-mean concentration trajectory for each demand-curve session.

### 4.10 Pose estimation and kinematic analysis

Behavioral videos were manually annotated using a custom-built Python graphical interface (Python 3.10, OpenCV 4.8, Tkinter) designed for pose annotation. A custom annotation interface was used to ensure precise control over keypoint definitions, consistent rat-centered cropping in low-light operant chambers, tight alignment with trial structure, and curated sampling of behaviorally relevant frames (e.g., stretch-attend, approach, withholding), producing cleaner training data and more reliable kinematic extraction than would be obtained with generalized pose-tracking frameworks. For each video frame, seven anatomical keypoints were labeled: head center, body center, tail, front right paw, front left paw, left back paw, and right back paw. The annotation tool provided frame-by-frame navigation, drag-and-drop point placement, interpolation between labeled frames, and quality control features including frame-by-frame review and correction capabilities. Annotations were exported in YOLO pose format, with each labeled frame producing: (1) a cropped image containing the rat, and (2) a corresponding text file specifying normalized bounding box coordinates (x_center, y_center, width, height) and keypoint positions (x, y, visibility) for all seven anatomical landmarks. Pose estimation models were trained using the YOLOv8n-pose architecture (Ultralytics YOLOv8) initialized from pretrained COCO-pose weights. The training dataset was split 80/20 into training and validation sets. Models were trained for 5 epochs with early stopping (patience = 3 epochs) using the following hyperparameters: image size = 640×640 pixels, batch size = 16, learning rate = 0.001 (with cosine decay), augmentation enabled (rotation *±*10°, translation *±*10%, scale *±*10%). Model performance was evaluated using mean average precision at IoU thresholds of 0.5 (mAP50) and 0.5:0.9 (mAP50-95) for both bounding box and keypoint detection. Final model selection was based on validation set mAP50-95 for keypoints. The best-performing model achieved mAP50 = 0.988 and mAP50-95 = 0.914 on the held-out validation set. Trained models were applied to all behavioral videos to generate frame-by-frame predictions of the seven keypoints. For each detection, the model returned (x, y) coordinates and confidence scores (0-1) for each keypoint. Predictions were retained only if: (1) detection confidence exceeded 0.5 for the bounding box, and (2) all seven keypoint confidence scores exceeded 0.5.

Trial-level movement features were extracted from tracked keypoint trajectories aligned to behavioral event timestamps from the operant system. To test whether gross motor output changed across the imaged sessions, we computed a session-level locomotion measure independent of trial structure. For each session, total locomotion was defined as the cumulative path length of the body-center keypoint via the sum of frame-to-frame Euclidean displacements over all retained frames of the session (pixels; chamber and camera geometry were held constant across the compared sessions, so pixel units are directly comparable across them). Frames failing the confidence criteria above were linearly interpolated before summation so that detection gaps did not inflate displacement. Total locomotion was compared across sessions with a one-way repeated-measures ANOVA (within-subject factor: session; *n* = 6 rats).

### 4.11 Surgical preparation and chronic miniscope imaging

Fourteen days before behavior, rats (n = 3 male, n = 3 female) were anesthetized with 1–3% isoflurane and placed in a custom robotic stereotaxic instrument [55]. The skull was leveled to bregma–lambda within *±*0.05 mm. A small craniotomy was made above right dorsomedial striatum (DMS). Animals received a 1,000-nL injection of AAV1-EF1*α*-jGCaMP8f-WPRE [56] (Addgene #176756) at 100 nL/min via a 33-gauge needle (NF33BV-2, WPI). Coordinates: AP -0.3 mm, ML +2.4 mm, DV -4.5 mm. After 10 min of diffusion, the needle was withdrawn. A GRIN lens (1 mm diameter, 9 mm length; GRINtech) was lowered to DV -4.4 mm and secured with jeweler’s screws, cyanoacrylate, and dental acrylic (Parkell C&B Metabond).

Animals received meloxicam (2.5 mg/kg s.c.) post-operatively and were monitored for *≥* 3 days. Calcium imaging was conducted using the Miniscope V4 (UCLA; www.miniscope.org) where, prior to the demand-curve start, animals underwent two dedicated habituation sessions on consecutive days in which they were connected to the miniscope and allowed to freely explore the chamber, ensuring acclimation to the head-mounted setup and minimizing handling- or equipment-related novelty during imaging days. The miniscope was connected via lightweight coaxial cable to the data acquisition system (Miniscope DAQ v3.2); video streams were saved as .avi with synchronized behavioral timestamps. A custom system for cable protection was used, consisting of (1) strain-relief epoxy at the baseplate–cable interface and (2) a flexible wire cage surrounding the coaxial cable from headstage to commutator. This was essential for recordings during high-motion and aversive-stimulation sessions.

### 4.12 Miniscope Processing Suite (MPS) pipeline

All calcium imaging data were processed using the Miniscope Processing Suite [18], our open-source GUI-based pipeline optimized for long-duration one-photon imaging. Processing parameters were held consistent across animals, with automatic parameter suggestions used to optimize performance for individual datasets. The pipeline performs preprocessing (background subtraction, spatial filtering, rigid translational motion correction, automatic detection and removal of erroneous frames), interactive field-of-view cropping, NNDSVD-based dimensionality reduction, watershed-based cell detection and initialization, constrained non-negative matrix factorization (CNMF) with autoregressive calcium dynamics for temporal refinement, dilation-bounded localized LASSO regression for spatial refinement, and a final temporal convergence pass with quality filtering on size, signal-to-noise ratio, and spatial-temporal correlation. Processed datasets were exported as compressed npy files containing spatial footprints (A), temporal components (C), deconvolved spike estimates (S), and background terms (b, f) with full metadata. All spatial footprints and calcium traces were manually inspected using the MPS Data Explorer interface and excluded if they exhibited vascular morphology, footprints outside the field of view, non-stationary baseline dynamics inconsistent with calcium transients, or anatomical localization outside the dorsomedial striatum based on histological verification. Full pipeline parameters and component-by-component validation are reported in [18]. Additionally, per-session yield varied across animals (one-way ANOVA, *F* (5, 30) = 4.22, *p* = 0.005; *η*^2^ = 0.41), consistent with expected animal-to-animal differences in viral expression and imaging-field placement, but was stable across sessions within animal, showing no systematic drift over the imaging window (Friedman *χ*^2^(5) = 6.42, *p* = 0.27; Spearman *ρ* = *−*0.12, *p* = 0.47).

### 4.13 Histology and viral injection confirmation

Following the conclusion of behavioral testing, animals were deeply anesthetized and transcardially perfused with phosphate-buffered saline followed by 4% paraformaldehyde. Brains were extracted, post-fixed overnight, and cryoprotected in 30% sucrose. Coronal sections (40 *µ*m) through the dorsomedial striatum were collected on a freezing microtome and immunostained for jGCaMP8f expression, with lens placement verified from the implant track. Sections were incubated with primary antibody chicken anti-GFP (1:1000, Aves Labs Cat#GFP-1020, RRID:AB_10000240) and detected with secondary antibody Alexa Fluor 488–conjugated goat anti-chicken IgY (1:500, Thermo Fisher Scientific Cat#A-11039, RRID:AB_2534096).

Sections were counterstained with DAPI, mounted, and imaged on a Keyence fluorescence microscope. Animals with viral expression or lens placement outside the dorsomedial striatum were excluded from analysis.

### 4.14 Demand-curve fitting and derived behavioral-economics parameters

Per-session fentanyl consumption was expressed as total intake (*µ*g/kg), computed as the number of reinforced deliveries in a session multiplied by the per-delivery dose defined above. For each animal, intake was averaged within each demand-curve price; the resulting per-animal values were then averaged across animals to give group consumption (*±* SEM) at each price. Vehicle sessions (0 *µ*g/mL) were summarized separately and excluded from the fit, as zero-dose consumption is undefined on the logarithmic price axis. Each fentanyl concentration was assigned a standardized unit price *C* (222.0, 125.0, 70.0, 40.0, and 22.0 *µ*g/mL mapped to *C* = 1, 5, 10, 25, 50, respectively), following the response-cost schedule and price assignments of Coffey et al. [30], with the highest dose anchored at *C* = 1 [cost convention of 30]. Group consumption was fit with the exponentiated demand model [57],

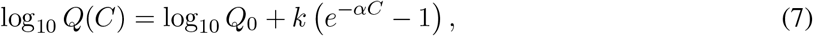

where *Q* is consumption (*µ*g/kg), *Q*_0_ is demand intensity (consumption as price approaches zero), *α* is the demand elasticity / essential-value parameter (smaller *α* = more inelastic demand), and *k* is the span of consumption in log_10_ units. Here *α* and *k* are the demand-model elasticity and span and are distinct from the pharmacokinetic rate constants *α* and *β* above and from the KNN parameter *k* in the S2C algorithm. *Q*_0_ and *α* were estimated by nonlinear least squares (`scipy.optimize.curve_fit`, bounded), with *k* fixed at 2.0 to stabilize the fit given the limited number of price points [58]; goodness of fit was summarized as *R*^2^. From the fitted curve we derived the standard behavioral-economics indices: maximal response output (*O*_max_) and the price of maximal output (P_max_) were taken as the maximum and the argmax, respectively, of the expenditure function *Q*(*C*) *· C* evaluated on a dense logarithmically spaced price grid, and essential value was computed as EV = 1*/*(100 *αk*^3*/*2^).

### 4.15 Cross-session DMS dynamics analysis

Press-aligned dynamics were computed from the event-aligned traces of Fig. 5d in a *−*2 to +10 s window around rewarded-press onset. Each neuron’s windowed trace was *z*-scored to its within-window mean and standard deviation, with the standard deviation floored at 0.5 so near-silent neurons could not dominate the normalization. The Σ Trace is the mean *z±* SEM across all segmented neurons that session; the heatmaps show the 100 neurons with the largest peak |*z*|, ordered by the temporal center of mass of their positive *z*-activity. Neurons are selected and ordered independently per session, so rows index population structure rather than tracked-cell identity across concentrations.

For the group-level quantification (Fig. 6e), neurons from all animals were pooled per session, the population peak time was taken as the Σ Trace maximum within 0–2 s of the press, and each neuron’s *z* at that time was summarized as the mean *±* SEM across the pooled population (*n* neurons); concentrations were compared by Kruskal–Wallis test with trained-vs-each Mann–Whitney *U* post hoc tests, treating each per-session population as an independent sample.

Single neurons within each concentration were partitioned by the shape of their press-aligned response (Fig. 6f, g). Each neuron’s per-neuron *z*-scored trace (computed as above) was restricted to the 0–10 s post-press window, reduced to its first 10 principal components, and clustered by Ward (variance-minimizing) agglomerative linkage; because the clustering input is per-neuron *z*, neurons are grouped by response *shape* rather than amplitude, which renders the partition gain-robust and licenses pooling neurons across animals within a concentration. Clusters were ordered by mean *z* in the 0–2 s press window, so that cluster 1 denotes the most press-excited type. The cluster count was fixed at *k* = 2 on the basis of a per-concentration model-selection sweep over *k* = 1–7, scored both by Gaussian-mixture Bayesian information criterion (BIC) and by mean silhouette: *k* = 1 was rejected at every concentration (ΔBIC = 511–858 for *k*=1 → 2), silhouette was maximal at *k* = 2 in every concentration (0.23–0.33) and never exceeded that value for *k ≥* 3, and although BIC continued to decrease weakly for larger *k* its minimizer was inconsistent across concentrations (*k* = 4–7) with only small gains (ΔBIC *≤* 243 relative to *k* = 2). Therefore, we implemented the best-separated two-type partition for further analyses. Clustering was performed independently per session on neurons pooled across all six animals and the representative-animal clusters in Fig. 6f, g illustrate the typology.

The two-type distinction was operationalized as a single fixed, dose-neutral criterion rather than the per-session Ward labels. A neuron was classed press-responsive if the peak of its per-neuron *z*-scored trace within 0–3 s of the rewarded press exceeded 2, and non-responsive. We took the fraction of segmented neurons in each class, summarized these as the across-animal mean *±* SEM, and compared classes across concentration with a Friedman test (Kendall’s *W* effect size), treating animals as repeated-measures blocks; this same fixed criterion defines the response classes carried into the cross-session tracking analyses (Fig. 6h) .

For the session-to-session flow analysis (Fig. 6i), we assigned the neurons that S2C registered across both sessions a given pair (222→70, 70→22, 22→0), and assigned each to press-responsive or non-responsive at each concentration by the same fixed criterion used for panel h. Each concentration column is normalized to unit height so that band heights are class fractions and each ribbon is drawn with width proportional to the number of co-tracked neurons making a given class-to-class transition. Transitions were summarized as the fraction of pair-tracked neurons that changed class between adjacent concentrations.

For the cross-session responsive-class analyses (Fig. 6j), each neuron S2C-registered across a session pair was labeled press-responsive or non-responsive at each session by the fixed *z >* 2 criterion above. Agreement between the two labels were quantified by the adjusted Rand index (ARI) over the cells co-tracked in that pair, with 95% confidence intervals from 2,000 cell-level bootstrap resamples. Turnover was defined as the fraction of co-tracked cells changing class, and retention the fraction of session-*s* responsive cells still responsive at session-*s*^*′*^, compared against the marginal responsive rate (chance). To separate biological reorganization from registration error we computed a registration ceiling expected if responsive identity were perfectly stable and the only disagreements arose from S2C mismatches. This simulation was performed assigning each cell its reference-session label and then each session’s label was independently flipped with probability 1 *−* F1 *≈* 0.015. The panel j raster visualizes this contrast, with the “Theoretical Stability” raster generated by the same procedure and the “Observed” raster showing the measured labels, both restricted to cells tracked across all four shown sessions and sorted by first-session status. As a stimulus-held-fixed control, the ARI and turnover analyses were repeated on the constant-concentration (70 *µ*g/mL) chain of consecutive sessions. To confirm the lability reflected the press response rather than the binary threshold, we recomputed cross-session agreement on a noise-robust graded-response measure via the baseline-subtracted mean of the press-window trace. This computation carried no cross-session structure specific to the press window, confirming the turnover is a property of the population and not an artifact of the binary criterion.

### 4.16 Statistical analysis

All analyses in Python 3.10 (SciPy, statsmodels). For the S2C-vs-ROI-matching comparison, the unit of paired analysis was a single F1 score per (animal, seed) within each (condition, neuron-count) cell, computed by averaging F1 across the four consecutive session transitions of that animal-seed’s 5-session chain. Each such F1 was paired across tools (S2C vs. CellReg) on identical underlying centroid data and ground-truth identity. The full paired-run count therefore arrives at 32 productive (condition *×* neuron-count) cells *×* 8 animals *×* 5 seeds = 1,280 paired comparisons in principle per footprint scenario. A paired comparison was excluded only when CellReg returned no defined F1 for any of the four consecutive transitions (Search-radius choice); with no per-pair F1 to pair against the corresponding S2C value, those runs cannot enter the paired statistic without imputation, which we declined to perform. This yields *n*_grand pooled_ = 1,262 in the worst-case scenario (18 dropped) and *n*_grand pooled_ = 1,262 in the best-case scenario (15 dropped). We note that this 1,262 is the unit of *statistical comparison*, not the count of S2C invocations: each chain involved 5 separate S2C runs (one per session) and produced 4 session-pair F1 values, so the underlying number of individual S2C runs is 5*×* larger (*∼*6,310) and the underlying number of session-pair F1 measurements is 4*×* larger (*∼*5,048). Per-condition tests used the paired Wilcoxon signed-rank statistic with Holm-Bonferroni correction across the 8 conditions within each neuron-count tier; effect sizes were reported as paired Cohen’s *d*_*z*_; 95% confidence intervals on the F1 difference were obtained via bias-corrected bootstrap (10,000 resamples). Pooled tests across conditions and neuron counts used the same paired-test framework on the full pooled set. For the temporal-correlation control, mean accuracy per duration was tested against chance (1*/N* = 0.10%) by paired *t*-test across the 10 session pairs. *α* = 0.05 throughout. Data are presented as mean *±* SEM unless otherwise noted.

## Data availability

The processed data supporting the findings of this study are deposited on Zenodo (DOI: 10.5281/zenodo.20938743) and released under CC-BY-4.0. This deposit is comprised of the MPS-extracted spatial footprints + traces, behavioral event tables, and S2C track assignments. The benchmark is reproducible from the released generator (Code availability). Raw single-photon miniscope videos (∼1.4 TB) are available from the corresponding author on reasonable request; they are not deposited due to their size and are not required to interpret, verify, or extend the reported results.

## Code availability

Stars2Cells is distributed as a GUI-driven standalone application for macOS (a signed and notarized DMG containing Stars2Cells.app) and Windows (a ZIP archive containing a standalone Stars2Cells.exe); both builds bundle a self-contained Python runtime, so users do not need to install Python or manage virtual environments. Source code is available at https://github.com/ariasarch/Stars2Cells_GUI, and the exact release used to produce the results in this manuscript is archived on Zenodo (DOI: 10.5281/zenodo.20938743). All other custom code used to generate the results reported here is collected in a single companion repository (https://github.com/ariasarch/S2C_COMPANION_REPO), archived on Zenodo (DOI: 10.5281/zenodo.20938869) All code is released under the MIT license.

## Acknowledgments

This work was supported by the National Institute on Drug Abuse (R01DA052618 and R00DA052571), a VA Puget Sound Health Care System Research & Development Pilot award, National Institute on Drug Abuse training grant T32DA007278.

## Author contributions

**A.P.-A**. Conceptualization; Methodology; Software; Validation; Formal analysis; Investigation; Data curation; Writing – original draft; Writing – review & editing; Visualization. **L.H**. Data curation (DMS dataset processing through MPS). **J.B**. Investigation (movement/kinematic analysis). **E.A**. Software (Astro.py porting); Validation (astrometric testing). **J.Q**. Investigation (DMS behavioral-economics fentanyl self-administration experiments, including animal training and histological verification). **K.R.C**. Conceptualization; Resources; Writing – review & editing. **J.F.N**. Conceptualization; Supervision; Funding acquisition; Resources; Project administration; Writing – review & editing.

## Competing interests

The authors declare no competing interests.

## Supplementary Information

### Supplementary Note 1: Ownership Guard Uniqueness

**Proposition 1** (Unique Emission). *Let Q* = *{P*_*a*_, *P*_*b*_, *P*_*c*_, *P*_*d*_*} be a quad with six pairwise distances. Under the ownership guard, Q is emitted exactly once across the entire diagonal set: from the single diagonal equal to its longest pairwise edge*.

*Proof*. Define the ownership diagonal of *Q* as the pair (*i*^*\**^, *j*^*\**^) achieving the maximum pairwise distance:

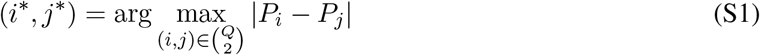

When processing diagonal (*d*_1_, *d*_2_), the ownership guard emits *Q* only if:

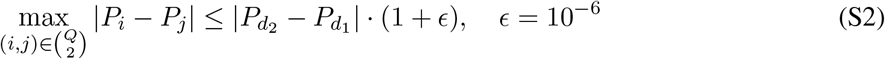

This condition is satisfied if and only if (*d*_1_, *d*_2_) *is* the longest edge (within floating-point tolerance). For generic point configurations (no two pairwise distances exactly equal), exactly one of the six edges is longest, so *Q* is emitted exactly once. Exact ties occur with probability zero for real-valued centroid coordinates.

#### Complexity reduction

Without the ownership guard, each quad could be emitted from up to 6 diagonals (one per pairwise edge that appears in the sampled diagonal set), requiring an *O*(*Q* log *Q*) global deduplication pass via np.unique. With the guard, the output is duplicate-free by construction and the deduplication pass is eliminated entirely.

### Supplementary Note 2: 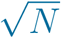 Threshold Scaling

The optimal descriptor similarity threshold *τ* must balance two forces: too low and same-cell matches are rejected (low recall); too high and false matches dominate (low precision). The per-animal calibration in Step 1.5 finds the optimum *τ* at each *N* by sweeping thresholds on sampled quad subsets, and the empirical scaling across the four neuron-count tiers tested is well-fit by 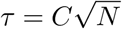. Below we derive this scaling from the mean-KNN-distance behavior of the local-diagonal class, and show why the alternative scaling implied by 4-D descriptor density is non-binding in the operating scenario.

#### Setup

S2C builds quads from two diagonal classes: (i) KNN-local diagonals connecting each neuron to its *k* nearest neighbors, and (ii) random long-range diagonals spanning the FOV. The two classes have very different diagonal-length scaling with *N* and, consequently, very different descriptor-blur behavior. The threshold is set by whichever class is worse-blurred, which turns out to be the local class.

**Step 1: local diagonal length scales as** *N*^*−*1*/*2^. For *N* points distributed in a fixed FOV of area *A*, the mean distance to the *k*-th nearest neighbor scales as

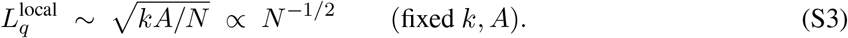

This holds for both Poisson and Poisson-disk-sampled point patterns; only the prefactor differs (Poisson-disk is more concentrated around its mean than Poisson, but both inherit the *N*^*−*1*/*2^ exponent from constant-density scaling). The intuition is direct: in a denser field, your nearest neighbor is simply closer.

**Step 2: same-cell descriptor blur scales as** 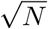 . Per-neuron centroid jitter of standard deviation *σ*_*j*_ propagates to descriptor space as 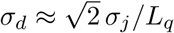 (Methods). Substituting Step 1:

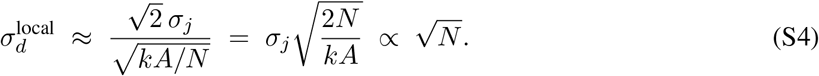

The intuition: at high density the same absolute jitter occupies a larger fraction of the (now shorter) diagonal length, and that ratio is exactly what propagates into descriptor space.

**Step 3: the threshold tracks same-cell blur**. A same-cell quad pair has descriptor distance *∼ σ*_*d*_ in expectation. For same-cell matches to survive descriptor filtering, *τ* must exceed *σ*_*d*_ by a constant capture margin *β >* 1:

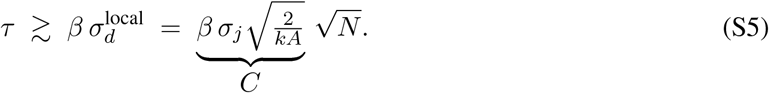

The constant *C* absorbs the capture margin *β* (set by the desired same-cell capture fraction; e.g., *β ≈* 2 captures *∼*95% of a Gaussian), the per-animal jitter level *σ*_*j*_, the usable FOV area *A*, the KNN parameter *k*, and an *O*(1) geometric prefactor relating mean KNN distance to typical local-quad diagonal length. None of these depend on *N*, so the leading-order behavior is exactly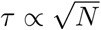. The empirical per-animal calibration of *C* recovers exactly this group of experimental constants.

#### The 4-D descriptor-density floor is non-binding

An alternative starting point is descriptor-space density: *Q ∝ N* quads occupy a compact 4-D region, so the expected nearest-neighbor distance among descriptors scales as NN_*d*_ *∝ N*^*−*1*/*4^ and the false-match floor in cosine distance shrinks accordingly. This argument predicts *τ* should also *shrink* with *N* – the opposite direction from both the empirical fit and Step 3 above. The reconciliation is that same-cell blur grows with *N* much faster than the false-match floor shrinks; their ratio is

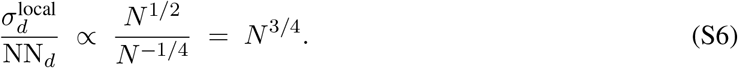

Throughout the operating range, same-cell blur is the binding constraint by a margin that widens with *N* ; the false-match floor is non-binding. False matches that survive the descriptor stage are filtered downstream by the consistency check and RANSAC, which are explicitly designed to absorb high false-match fractions.

#### Why long-range diagonals are insensitive to *N*

Random long-range diagonals span the FOV regardless of density, 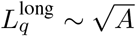, so their descriptor blur 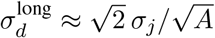is independent of *N* . This is the design reason both classes exist. Local diagonals stabilize matching under small inter-session shifts but set the threshold; long-range diagonals carry the matching signal at high density and operate well below *τ*. The two-class construction is what makes per-animal calibration tractable: *C* only needs to track local-diagonal blur, while the bulk of useful matches come from the long-range class.

### Supplementary Note 3: Coverage Remediation

The ownership guard (Supplementary Note 1) eliminates duplicate quads but can cause neurons in dense local regions to have zero or very few quads. This occurs when a neuron only participates in short diagonals, all of which have their quads claimed by longer neighboring diagonals.

#### Detection

For each neuron i, define coverage κ_*i*_ as the number of quads containing *i*. The field median coverage is 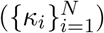. Neuron *i* is undercovered if:

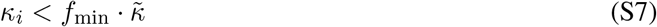

where *f*_min_ = 0.4 by default.

#### Remediation

For each undercovered neuron *i*, all diagonals (*i, j*) and (*j, i*) in the diagonal bucket dictionary are re-processed by _process_diagonal with enforce_ownership=False. The resulting quads are subject to all other quality filters (area *≥* 1.0, min pairwise distance, non-degeneracy) and deduplicated against the existing quad set via a Python set of sorted index tuples. Quality pruning (quad_keep_fraction) is applied to the remediation batch independently.

#### Guarantees

Remediation quads may duplicate quads that would have been emitted from a different diagonal under normal ownership. However, because they are deduplicated against the existing set, no quad appears twice in the final output. The trade-off is that these remediation quads lose the automatic “emitted exactly once” guarantee the ownership guard gives the main pass: because they are produced by diagonals that are not their rightful owner, their uniqueness is enforced by the explicit duplicate check rather than guaranteed by construction.

### Supplementary Note 4: Computational Complexity

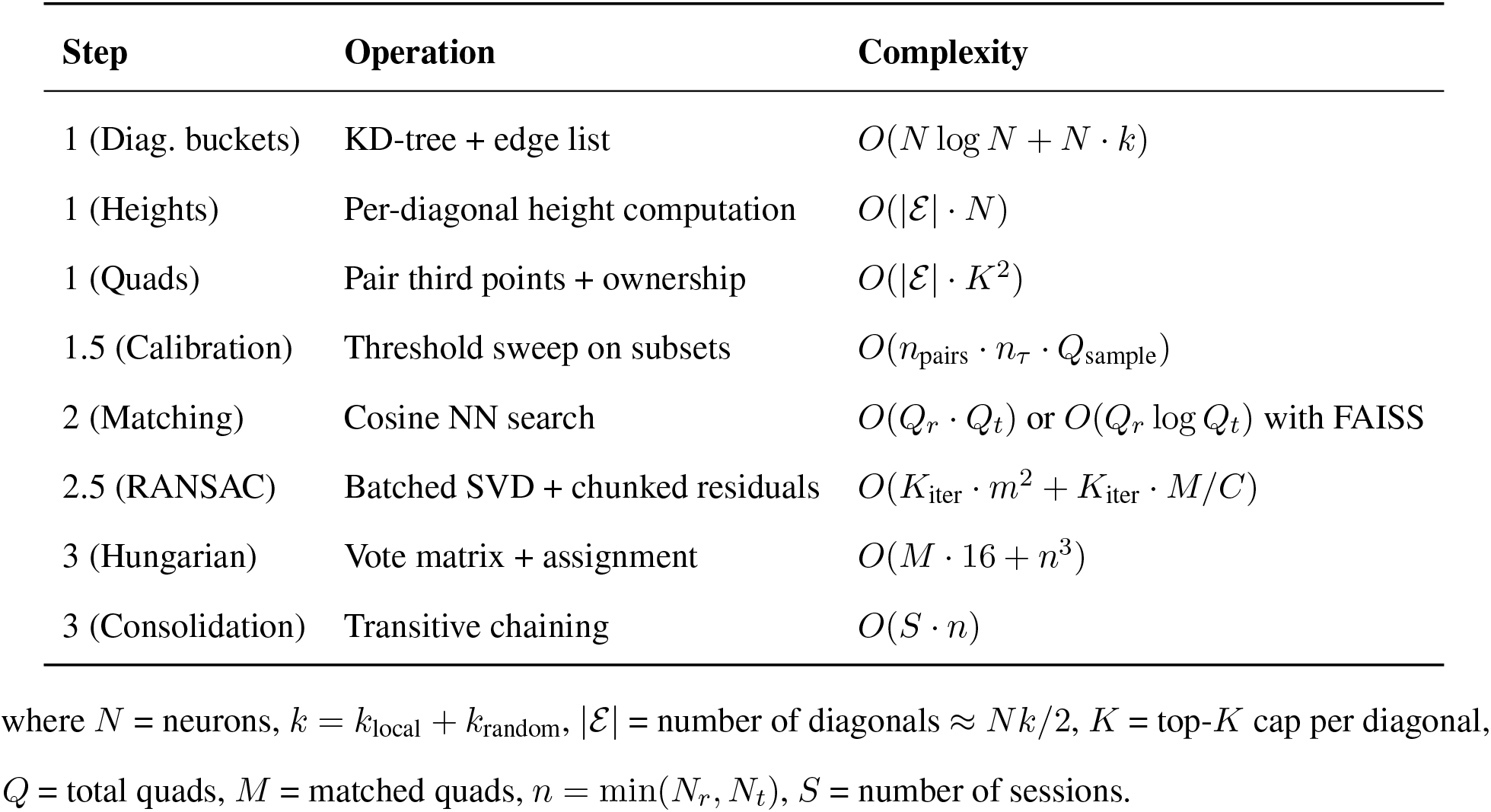

### Supplementary Note 5: Synthetic Degradation Model

#### Base field generation

For each synthetic animal, a base centroid field 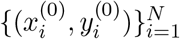 is placed in a 600 *×* 600-pixel FOV by Bridson Poisson-disk sampling, producing a blue-noise (well-separated, non-clustered) layout. The minimum spacing *r*_min_ defaults to 55% of the theoretical hex-packing maximum for the requested *N* within the usable FOV (margin = 10 px), so density automatically scales with neuron count. A fallback rejection-sampling step with relaxed *r*_min_ (factor 0.8) handles edge cases where Bridson cannot place all requested points in the first pass.

#### Per-neuron base jitter

Independent of session-to-session perturbations, each neuron carries a stable per-neuron centroid offset 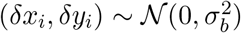 with *σ*_*b*_ = 0.5 px, modeling systematic CNMF centroid bias that persists across sessions for that neuron. This is added in addition to a per-session perturbation *σ*_*p*_ = 0.2 px drawn fresh each session.

#### Session generation

For session *s ∈ {*1, …, *S}* (where *S* = 5), the base field is transformed in the following fixed order:

1. **Rotation:**, 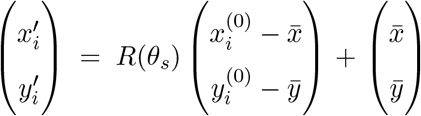where 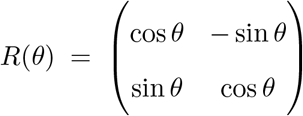 with *θ*_*s*_ accumulating across sessions in Tiers A and B and following the random-walk schedule in Tier C.
2. **Translation:** 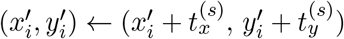, similarly accumulating per tier.
3. **Clip-to-FOV:** any neuron whose transformed position falls outside [0, 600]^2^ is removed, modeling loss off the sensor edge after large transforms (this is what makes large rotation/translation conditions genuinely hard rather than merely shifted).
4. **Dropout (sessions** *≥* 2 **only):** a fraction *δ*_*s*_ of remaining neurons is randomly removed, modeling CNMF detection failure. *No new neurons are spawned* – this matches the biological fact that ROIs can fail to be detected in a given session, but new cells do not spontaneously appear.
5. **Jitter:** 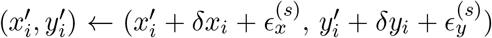, where 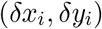 is the stable per-neuron base offset and 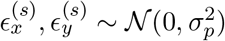 is the per-session perturbation, applied only to neurons that survived clipping and dropout.
6. **Ground-truth capture:** the surviving base indices (before permutation) are stored as ground_truth _base_ids in the per-file ground-truth dictionary.
7. **ID permutation:** ROI identifiers are randomly shuffled to prevent matching by index.

### Supplementary Note 6: CellReg Reimplementation and Validation

#### ROI-based matching via a probabilistic model

CellReg [9] models the probability that neurons *i* (session 1) and *j* (session 2) are the same cell as:

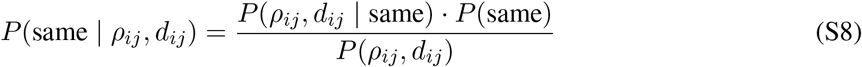

where *ρ*_*ij*_ is the spatial footprint correlation and *d*_*ij*_ is the centroid distance. The likelihoods are modeled as:

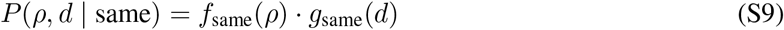

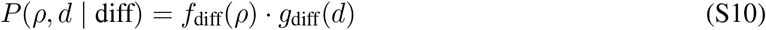

where *g*_same_(*d*) is modeled as a Rayleigh distribution (centroid distances for matched neurons) and *g*_diff_(*d*) as approximately uniform over the FOV. The correlation distributions *f*_same_(*ρ*) and *f*_diff_(*ρ*) are estimated non-parametrically from the data. Parameters are fit via expectation-maximization over the observed pairs.

#### Python reimplementation

The step-by-step port of CellReg’s spatial-correlation pathway, its cluster_cells final-assignment step (which differs from a one-shot Hungarian solve), and its pair-for-pair validation against native MATLAB CellReg are given in Methods (“ROI-based matching reimplementation”). The probabilistic model above is what those steps implement.

#### Parallelization

The validated reimplementation was wrapped in an embarrassingly-parallel batch driver (Python’s concurrent.futures.ProcessPoolExecutor, up to 50 workers, each pinned to a single BLAS thread) so the full benchmark runs on one workstation; the algorithm is unchanged and only its execution is parallelized.

### Supplementary Note 7: Full Per-Condition Benchmark Statistics

Table 1 expands the pooled and per-neuron-count summaries in the main text (Fig. 4g) to all 32 (condition *×* neuron-count) benchmark cells. S2C’s F1 advantage over ROI-based matching is positive and significant in every cell, with the smallest gaps on the mildest perturbation (Tier-A translation) and the largest on the high-displacement Tier-B/C conditions.

**Table 1:** Per-condition S2C vs. ROI-based matching (worst-case footprint scenario). Mean F1 per tool, the paired S2C *−* ROI F1 difference with 95% bootstrap CI, paired Cohen’s *d*_*z*_, and Wilcoxon *p* (Holm-corrected across conditions within each neuron-count tier). *n* = 40 paired runs per cell except where CellReg returned no defined F1 (excluded per Methods); S2C precision was *≥* 97.7% throughout; best-case footprint values are near-identical.

| Condition | $n$ | S2C F1 (%) | ROI F1 (%) | ΔF1 (pp) [95% CI] | $d_z$ | $p_{\text{Holm}}$ |
| --- | --- | --- | --- | --- | --- | --- |
| *N = 100 neurons* |  |  |  |  |  |  |
| A: rotation | 40 | 99.6 | 76.0 | +23.5 [+15.2, +32.4] | +0.82 | $2.3 \times 10^{-9}$ |
| A: translation | 40 | 99.4 | 97.8 | +1.6 [+0.8, +2.6] | +0.57 | $9.4 \times 10^{-4}$ |
| A: combined | 40 | 99.4 | 68.8 | +30.6 [+22.0, +39.3] | +1.10 | $1.5 \times 10^{-11}$ |
| B: dropout | 40 | 99.9 | 47.9 | +52.0 [+46.0, +57.6] | +2.68 | $1.5 \times 10^{-11}$ |
| B: drift | 35 | 99.8 | 19.3 | +80.6 [+74.9, +85.9] | +4.84 | $1.7 \times 10^{-10}$ |
| B: rotation | 39 | 99.9 | 14.4 | +85.5 [+79.9, +90.1] | +5.15 | $1.8 \times 10^{-11}$ |
| B: combined | 39 | 99.8 | 12.3 | +87.5 [+83.2, +91.2] | +6.75 | $1.8 \times 10^{-11}$ |
| C: random walk | 40 | 99.7 | 23.1 | +76.6 [+70.9, +82.0] | +4.17 | $1.5 \times 10^{-11}$ |
| *N = 250 neurons* |  |  |  |  |  |  |
| A: rotation | 40 | 99.9 | 69.0 | +30.9 [+21.5, +41.0] | +0.98 | $1.5 \times 10^{-11}$ |
| A: translation | 40 | 99.9 | 97.4 | +2.5 [+1.2, +4.0] | +0.54 | $6.3 \times 10^{-7}$ |
| A: combined | 40 | 99.9 | 62.0 | +37.9 [+28.7, +47.0] | +1.24 | $1.5 \times 10^{-11}$ |
| B: dropout | 40 | 99.9 | 36.8 | +63.1 [+59.1, +67.1] | +4.81 | $1.5 \times 10^{-11}$ |
| B: drift | 38 | 99.9 | 15.4 | +84.5 [+78.7, +89.9] | +4.86 | $1.6 \times 10^{-7}$ |
| B: rotation | 40 | 99.9 | 7.6 | +92.4 [+89.9, +94.5] | +12.27 | $1.5 \times 10^{-11}$ |
| B: combined | 39 | 99.9 | 6.8 | +93.1 [+91.2, +94.8] | +15.58 | $1.6 \times 10^{-7}$ |
| C: random walk | 40 | 99.9 | 20.1 | +79.8 [+76.5, +82.8] | +7.63 | $1.5 \times 10^{-11}$ |
| *N = 500 neurons* |  |  |  |  |  |  |
| A: rotation | 40 | 97.2 | 54.7 | +42.4 [+33.4, +51.3] | +1.45 | $1.6 \times 10^{-11}$ |
| A: translation | 40 | 97.4 | 92.9 | +4.5 [+2.3, +6.9] | +0.61 | $4.6 \times 10^{-3}$ |
| A: combined | 40 | 97.2 | 45.0 | +52.2 [+44.2, +59.7] | +2.07 | $1.5 \times 10^{-11}$ |
| B: dropout | 40 | 97.0 | 30.5 | +66.5 [+63.9, +69.0] | +7.86 | $1.5 \times 10^{-11}$ |
| B: drift | 35 | 97.0 | 11.3 | +85.6 [+81.5, +89.5] | +6.97 | $1.2 \times 10^{-10}$ |
| B: rotation | 40 | 96.7 | 6.3 | +90.4 [+88.1, +92.4] | +12.78 | $1.5 \times 10^{-11}$ |
| B: combined | 39 | 96.8 | 5.2 | +91.6 [+90.2, +92.9] | +21.03 | $1.5 \times 10^{-11}$ |
| C: random walk | 40 | 96.7 | 15.4 | +81.3 [+78.5, +83.9] | +9.19 | $1.5 \times 10^{-11}$ |
| *N = 1,000 neurons* |  |  |  |  |  |  |
| A: rotation | 40 | 94.3 | 39.8 | +54.4 [+45.9, +62.7] | +1.97 | $1.5 \times 10^{-11}$ |
| A: translation | 40 | 94.9 | 84.3 | +10.6 [+6.8, +14.6] | +0.82 | $2.6 \times 10^{-4}$ |
| A: combined | 40 | 94.2 | 32.7 | +61.5 [+55.0, +67.4] | +3.07 | $1.5 \times 10^{-11}$ |
| B: dropout | 40 | 99.3 | 23.2 | +76.1 [+75.0, +77.0] | +22.98 | $1.5 \times 10^{-11}$ |
| B: drift | 38 | 100.0 | 7.9 | +92.1 [+89.2, +94.8] | +10.12 | $2.2 \times 10^{-11}$ |
| B: rotation | 40 | 99.8 | 4.4 | +95.3 [+94.0, +96.6] | +22.20 | $1.5 \times 10^{-11}$ |
| B: combined | 40 | 97.4 | 3.4 | +93.9 [+92.0, +95.6] | +15.70 | $7.1 \times 10^{-8}$ |
| C: random walk | 40 | 98.4 | 10.5 | +87.9 [+85.9, +89.8] | +13.90 | $1.5 \times 10^{-11}$ |

## Notes

### Competing Interest Statement

The authors have declared no competing interest.

